# Inference of past demography, dormancy and self-fertilization rates from whole genome sequence data

**DOI:** 10.1101/701185

**Authors:** Thibaut Sellinger, Diala Abu Awad, Markus Möst, Aurélien Tellier

## Abstract

Several methods based on the Sequential Markovian Coalescent (SMC) have been developed to use full genome sequence data to uncover population demographic history, which is of interest in its own right and a key requirement to generate a null model for selection tests. While these methods can be applied to all possible species, the underlying assumptions are sexual reproduction at each generation and no overlap of generations. However, in many plant, invertebrate, fungi and other species, those assumptions are often violated due to different ecological and life history traits, such as self-fertilization or long term dormant structures (seed or egg-banking). We develop a novel SMC-based method to infer 1) the rates of seed/egg-bank and of self-fertilization, and 2) the populations’ past demographic history. Using simulated data sets, we demonstrate the accuracy of our method for a wide range of demographic scenarios and for sequence lengths from one to 30 Mb using four sampled genomes. Finally, we apply our method to a Swedish and a German population of *Arabidopsis thaliana* demonstrating a selfing rate of *ca.* 0.8 and the absence of any detectable seed-bank. In contrast, we show that the water flea *Daphnia pulex* exhibits a long lived egg-bank of three to 18 generations. In conclusion, we here present a novel method to infer accurate demographies and life-history traits for species with selfing and/or seed/egg-banks. Finally, we provide recommendations on the use of SMC-based methods for non-model organisms, highlighting the importance of the per site and the effective ratios of recombination over mutation.

## 1 Introduction

Genomes, and especially genetic polymorphism, are shaped by molecular forces, such as mutation and recombination, but also ecological forces intrinsic to, or independent of, the biology of the species [11]. Therefore, polymorphism data contain a plethora of information that goes beyond the physiological functions encoded therein. Recent advances in sequencing technologies enable us to obtain whole genome data for many individuals across several populations even for non-model species [49, 50, 25, 36]. In particular, inferring demography is of interest in its own right so as to understand the history of existing and/or extinct species (population expansion, colonization of new habitats, past bottlenecks) [58, 28, 36]. Inferring demography is also necessary to generate null models for outlier scans, *e.g.* scanning genomes for genes under selection. Indeed, wrongly estimated demographies may strongly bias the outcome of such scans [37]. It is now common practice to simulate the past demography of a population, as a null model, to set up thresholds for selection scan methods. Therefore, an accurate demographic inference should yield more reliable selection results [43, 37]. To this aim, new models and methods have been developed to extract previously unavailable information from whole genome sequence data [48, 22, 40, 41, 26]. Inference is based on modelling single nucleotide polymorphism (SNPs) along the genome across individuals, the density of which results from the interplay between mutation, time to ancestral common ancestors and recombination. Thus, the common denominator in all these methods is their reliance on the per site molecular ratio of the recombination (*r*) and the mutation (*μ*) rate of the species 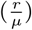, more precisely on its effective value 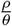. We classically define *ρ* as the effective recombination rate and *θ* as the effective mutation rate. We note that so far, applications of these approaches have considered these ratios as interchangeable [40, 48, 22], which is a strong assumption and may be violated in some species that do not fulfill the assumptions of the classic Wright-Fisher diploid model with two sexes (*e.g.* equal sex-ratio, sexual reproduction at each generation and no overlap of generations). In humans and mammals, as *ρ* = 4*N*_*e*_*r* and *θ* = 4*N*_*e*_*μ* (*N*_*e*_ being the effective population size), we indeed find 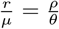. Yet, even in this case, bias can be found if 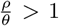[48]. In such cases, the number of mutations is not sufficient to detect all recombination events. The model is therefore no longer able to correctly estimate the Ancestral Recombination Graph, *i.e* the superposition of coalescent trees at different positions on the genome to display genealogies of sequences in the presence of recombination. Generally, it becomes necessary to extend existing approaches to account for characteristics and traits of species that can influence *ρ* or *θ*, and thus define when these methods can be accurately applied. It is additionally of interest to assess the accuracy of such methods for various values of the ratio 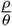.

Current methods rely on the Sequentially Markovian Coalescent (SMC) [30, 27] to account for the linear structure of genome sequences. SMC models the genealogy of a sample along a chromosome. First, a genealogy is built under the neutral coalescent and, in a second step, recombination and linkage disequilibrium are incorporated using a Poisson process [54, 55]. Applying Hidden Markov Models, it therefore becomes possible to calculate the probabilities of whole genome sequence data and infer the most likely values of the model parameters. These approaches can thus infer 1) the changes in population size (*χ*_*t*_, where 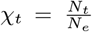, *N*_*e*_ and *N*_*t*_ being the effective population size and the current population size at time t, respectively) by inferring any variation of the coalescent rate in time, and 2) the ratio of effective recombination over the effective mutation rate 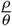. From this ratio, the recombination rate can be estimated if the mutation rate is known (assuming 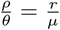).

The described methods have been almost exclusively built to be applied to hominid data, therefore rely on several assumptions that are violated in many species (and most likely also in hominids): non overlapping generations, equal sex ratio, sexual reproduction through outcrossing. Indeed, with the rise of next-generation sequencing technology, these methods are now frequently applied to whole genome sequences of species with characteristics that greatly differ from humans [15, 25, 9, 50]. In many species (*e.g.* plants, invertebrates) life-history strategies, such as mating systems or offspring production, influence the relationship between *r* and *ρ* and *μ* and *θ* [11]. If these effects are not accounted for, inferences using these methods may be biased and lead to misinterpretation of the results.

Two very common features in plant and invertebrate species are the maintenance of offspring as seed or egg-banks [6, 12, 5] and self-fertilization [17]. Indeed, as a consequence of environmental fluctuations, species can develop bet-hedging strategies such as seed-banking [45, 13, 20]. It increases the observed diversity [51, 33] and affects the rate of selection and neutral genomic evolution [16, 46]. Due to the discrepancy between census (*N*_*cs*_) and effective population size (*N*_*e*_) caused by seed-banks [47, 33], we expect that 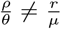. Seed-banks can therefore strongly bias demographic inference if ignored [59]. Self-fertilization, on the other hand, decreases the effective population size. This reproductive strategy has evolved many times independently, and is one of the most common evolutionary transitions observed in flowering plants [4]. The main consequence of this mating system is an increased homozygosity, which directly results in a decreased effective recombination rate (*ρ*) compared to the molecular recombination rate *r* (since recombination events between two homozygous haplotypes are invisible), as well as a reduction in genetic diversity [3]. Due to their contradictory effects on the effective population size, the simultaneous occurrence of these traits (dormancy and self-fertilization) may in fact be missed, and extensions of inference methods to account for them could not only allow for more accurate inferences of parameters and demographic histories of species with these traits, but could also provide a means with which to detect their respective rates.

To account for self-fertilization and seed-banks (or egg-banks) we develop a modified version of PSMC’ [40], named ecological Sequentially Markovian Coalescent (eSCM). Our model uses the deviation between the ratios 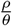 and 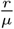 to infer self-fertilization and the existence of seed-banks. We first apply eSMC to simulated data to demonstrate its accuracy and then to genome sequence data of a plant, *Arabidopsis thaliana*, and an invertebrate species, *Daphnia pulex*. In both species self-fertilization and/or seed/egg-banks have been observed or suspected. *A. thaliana* presents a very high self-fertilization rate of 99% [1] and it has been suggested that populations in Scandinavia may have evolved seed-banks in Sweden [19] and Norway [24]. *D. pulex* exhibits cyclical parthogenesis and is known to have a dormant stage in eggs produced through sexual reproduction [25, 10]. These resting eggs possibly can potentially build up an egg-bank in the lake sediment as observed in many *Daphnia* species [6, 2].The aim of our study is thus three-fold. First we build a method based on the SMC using polymorphism data to infer the germination and/or self-fertilization rates jointly with the past demographic history. Second, we study the effect of variable ratios of 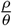 (and 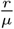) on the accuracy of past demographic estimates. Third, we apply our of method to existing datasets from *Arabidopsis thaliana* and *Daphnia pulex* which have well documented high self-fertilization rates and egg-banks, respectively. We find a strong signature of self-fertilization in *Arabidopsis thaliana* and a strong signature of egg-banks in *Daphnia pulex*. We find that self-fertilization has little effect on the inference of the demographic history, whereas neglecting seed-banks can strongly affect the inferred population size.

## 2 Overview of the model

### 2.1 The coalescent with seed-bank and self-fertilization

We model population seed-banks using the same hypotheses described in [18]. Under these assumptions, seed-banking can be accounted for by rescaling the coalescent rate by *β*^2^, where *β* (0 ≤ *β* ≤ 1) is the germination rate, or the expected germination probability at every generation (*β* = 1 implying that there is no seed-bank). The probability that two lineages find a common ancestor in the active population size is slowed by a factor *β* × *β* in the active population when looking backward in time. Hence, the expected coalescent times are therefore increased by a factor 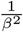. Assuming mutations can arise during the dormant stage at the same rate as in the active population, we expect to have 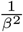 more mutations [33, 45, 16]. However, because recombination only occurs in the active population and concerns only one lineage backward in time, it is rescaled by *β* [46]. Therefore, although coalescent times are 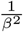 longer, there are only 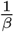 more expected recombination events, giving the relationship 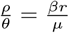.

To model self-fertilization, we adopted the island model described in [32], where *σ* (0 ≤ *σ* ≤ 1) represents the proportion of offspring produced through self-fertilization (if *σ* = 1 all individuals are produced through self-fertilization). As a consequence, the coalescent rate is increased by a factor 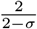 [31] and the recombination rate is decreased by a factor 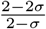 [32] since recombination events in homozygous individual are invisible. In the case of self-fertilization, we thus find the following relationship: 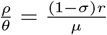.

To simultaneously model seed-banking and self-fertilization we assume their effects to be independent and that there is no correlation between dormancy and the rate of self-fertilization. Under this assumption we can simply multiply their effects as in [51], giving the relationship 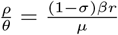. We therefore have a potential confounding effect between self-fertilization and seed-banking when observing the recombination and mutation ratio 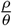. Because of their opposing effects on the effective population size (seed dormancy increasing it, and self-fertilization decreasing it), the effects of these traits can be compensated by one another. As consequence, in our model seed-banking can theoretically be equivalent to self-fertilization with higher effective population size (see the discussion for ruling this effect out in practice).

### 2.2 ecological Sequentially Markovian Coalescent (eSMC)

eSMC is an extension to the PSMC’ algorithm [40] and is therefore a Hidden Markov Model (HMM) along two haplotypes. It adds the possibility to account for and estimate germination rates *β* (seed and egg-banks), as well as self-fertilization rates *σ*, while inferring the demographic history of a population/species (*χ*_*t*_). The input file contains the comparison of two genome sequences; at each position if the two nucleotides on each sequence are the same, this is indicated by 0, otherwise by 1.

The general idea is that mutations are more likely to arise if the coalescent time is long between the two lineages, therefore, as in PSMC’, the hidden states are coalescent times (where time is discretized). In SMC, the coalescent model is the classic n-Kingman coalescent with sample of size two with the assumption of the classic Wright-Fisher model. However, because seed-banks and self-fertilization scale the coalescent rate, we re-scale the hidden states by 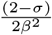, where *σ* and *β* are respectively the self-fertilization rate and the germination rate. Variations in the coalescent times, or the hidden states, along the sequence, are caused by recombination events. In order to account for the effects of seed-banks and self-fertilization on the observed variations, the recombination rate must be re-scaled by 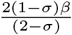. A detailed description of the model can be found in section 1 of the appendix.

We use a Baum-Welch Algorithm to find the estimates of our parameters (the effective recombination rate *ρ*, the germination rate *β*, the self-fertilization rate *σ*, and changes in population size *χ*_*t*_ [40, 48] (Table 1). The Baum-Welch Algorithm aims to maximize an objective function (*L*) analogous to the likelihood [40].

**Table 1:**
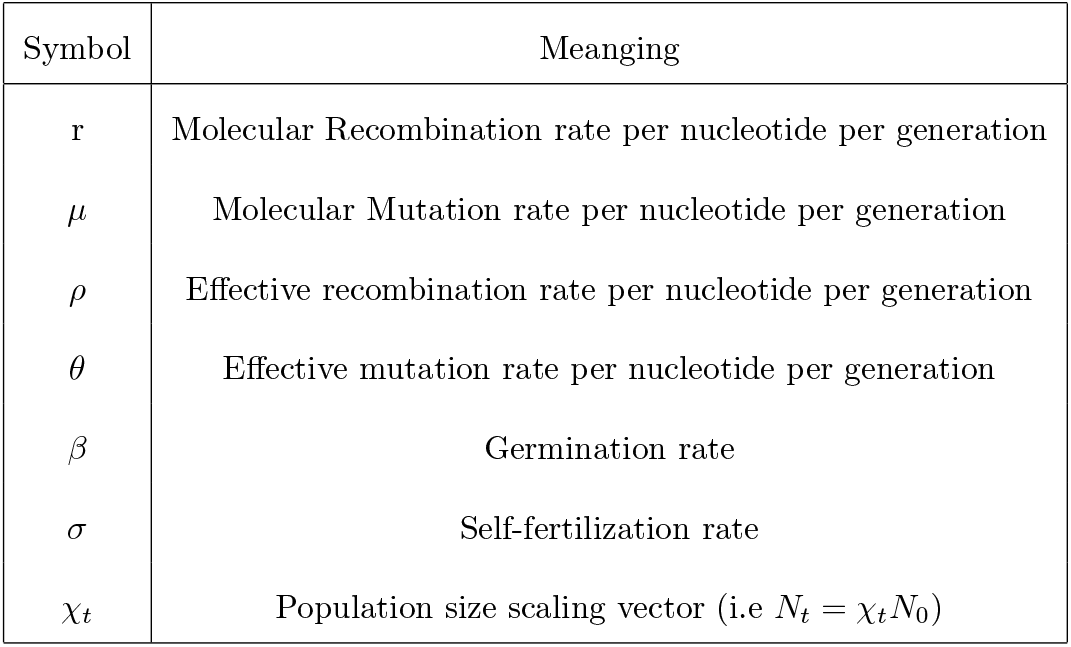
Symbol table

| Symbol | Meanging |
| --- | --- |
| $r$ | Molecular Recombination rate per nucleotide per generation |
| $\mu$ | Molecular Mutation rate per nucleotide per generation |
| $\rho$ | Effective recombination rate per nucleotide per generation |
| $\theta$ | Effective mutation rate per nucleotide per generation |
| $\beta$ | Germination rate |
| $\sigma$ | Self-fertilization rate |
| $\chi_t$ | Population size scaling vector (i.e $N_t = \chi_t N_0$) |

Finally, to increase the model’s accuracy, more than two sequences can be used. Assuming that every individual comes from the same population, we compute a composite objective function *CL*, which is the product of all the objective functions. Therefore if we have *n* phased sequences we have 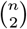 analyses. If sequences are not phased we have *n* analyses (one for each diploid individual). The algorithm is run to chiefly infer *β*, *σ* and *χ*_*t*_ by fixing or simultaneously inferring the ratio 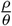 given an initial value of this ratio.

Our model is implemented in the R-package eSMC and can be downloaded from our GitHub repository (https://github.com/TPPSellinger/eSMC, note that the R package devtools is required for the installation). The algorithm uses the same input file format as MSMC or PSMC’. Note that in comparison with other similar models, our implementation in R is thus slightly slower, but still allows whole genome sequence analysis of several individuals in a reasonable amount of time. Furthermore, we added a feature to eSMC allowing the simulation of pseudo-observed data based on the obtained results. These pseudo-observed data are then analyzed by eSMC to asses the robustness of the results.

## 3 Results

We first study the theoretical accuracy and properties of our method on sequence data simulated under different scenarios. We then analyze real sequence data from two European populations of *Arabidopsis thaliana* (Sweden and Tübingen Germany), in which seed-banking is suspected in northern populations, while accounting for self-fertilization. We also analyze data from *Daphnia pulex*, for which egg-banks are known to be a prominent biological feature.

### 3.1 Theoretical results

#### 3.1.1 Convergence property in the absence of seed-banks and self-fertilization

We start by analyzing 4 sequences of 30 Mb from a population simulated under a “saw-tooth” scenario (repetitions of expansion followed by a decrease) without seed-banks or self-fertilization (Figure 1, similar to those used in [40, 48]). As previously known, with a population size of approximately 10^4^ inference of very recent (< 10^2^ generations) and very old events (> 10^5^ generations) cannot be recovered due to limits of the coalescent resolution. All of the methods tested with this scenario (eSMC, MSMC, MSMC2 and PSMC’) show an accurate fit and small variability of the estimated demo-graphic histories. Nonetheless, eSMC shows much less variability in more recent times compared to the other methods. Estimates of 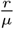 by eSMC, PSMC’ and MSMC are accurate (Table 2), whereas MSMC2 exhibits a strong bias, generally underestimating the recombination rate in this scenario.

**Figure 1:**
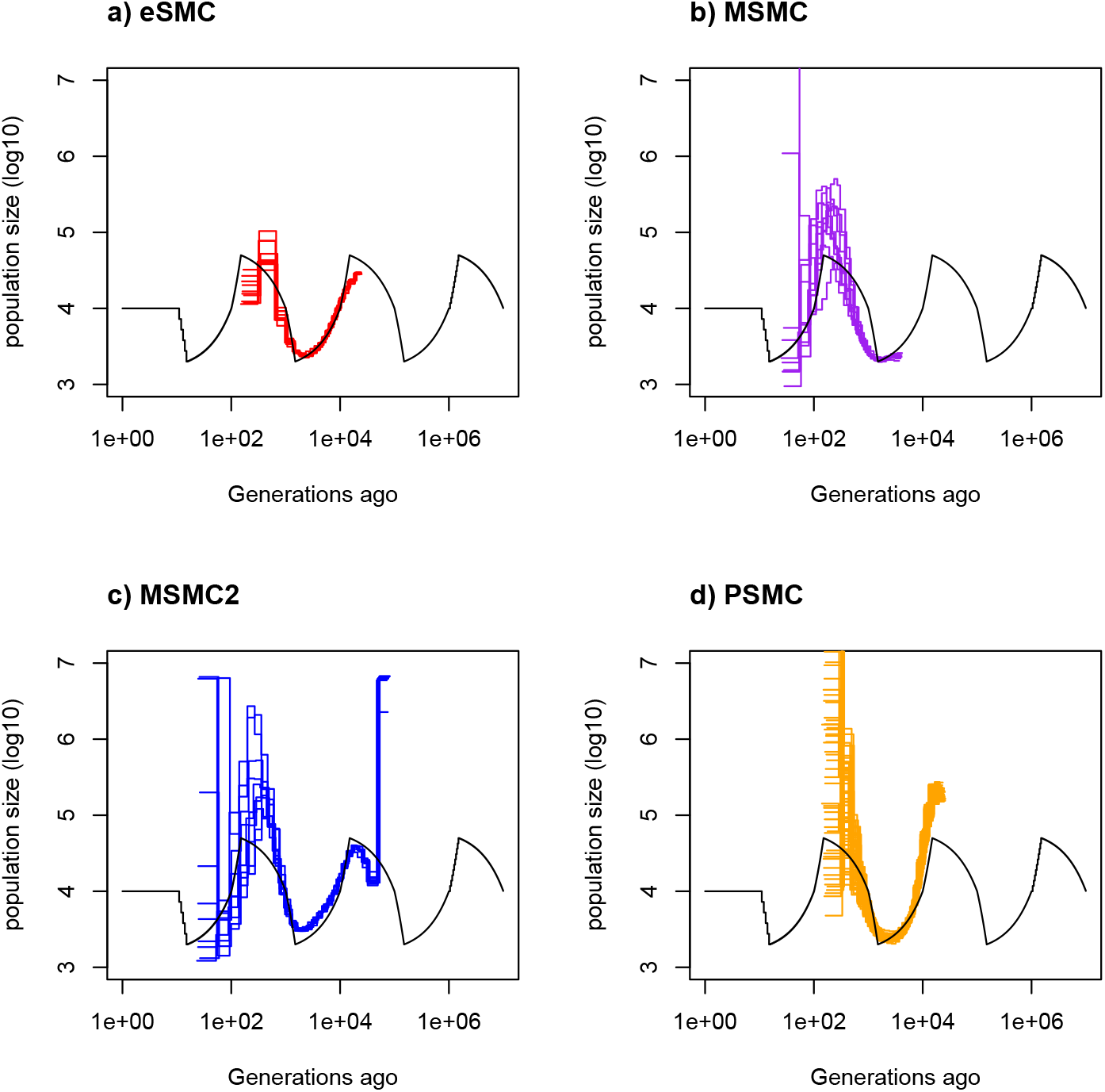
Estimated demographic history using four simulated sequences of 30 Mb under a saw-tooth scenario with 10 replicates. Mutation and recombination rates (respectively *μ* and *r*) are set to 2.5 × 10^−8^ per generation per bp. Therefore 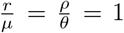. The simulated demographic history is represented in black. a) Demographic history estimated by eSMC (red). b) Demographic history estimated by MSMC (purple). c) Demographic history estimated by MSMC2 (blue). d) Demographic history estimated by PSMC’ (orange).

**Table 2:** Averaged estimated values for the recombination over mutation ratio 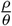 over ten repetitions. The “saw-tooth” estimations are seen in Figure 1, and the other scenarios are shown in Supplementary Figures 3-6.

| Scenario | Sequence length | real $\frac{\rho}{\theta}$ | eSMC $\frac{\rho}{\theta}$ | PSMC $\frac{\rho}{\theta}$ | MSMC $\frac{\rho}{\theta}$ | MSMC2 $\frac{\rho}{\theta}$ |
| --- | --- | --- | --- | --- | --- | --- |
| "Saw-tooth" | 30 Mb | 1 | 0.90 | 0.93 | 1.28 | 0.38 |
| "Saw-tooth" | 10 Mb | 1 | 0.94 | 0.96 | 1.30 | 0.39 |
| "Saw-tooth" | 10 Mb | 5 | 2.20 | 2.06 | 2.89 | 1.49 |
| "Saw-tooth" | 1 Mb | 1 | 0.96 | 1.00 | 1.31 | 0.86 |
| Constant | 10 Mb | 1 | 0.96 | 0.95 | 1.04 | 0.80 |
| Bottleneck | 10 Mb | 1 | 1.08 | 1.08 | 1.08 | 1.00 |
| Expansion | 10 Mb | 1 | 1.05 | 1.05 | 1.08 | 0.98 |
| Decrease | 10 Mb | 1 | 0.91 | 0.90 | 1.06 | 0.70 |

With shorter sequences of 10 MB (Supplementary Figure 1), eSMC and PSMC’ display convergence properties similar to those with 30 Mb sequences. However, MSMC and MSMC2 result in higher variance and less accurate estimation of the demographic histories. As eSMC has good convergence properties with 10 Mb sequences, all the following analyses are carried out with this sequence length, which is a more realistic length when considering real data. In addition, since the longest scaffolds from *Daphnia pulex* are in fact very short (length shorter than 2Mb) we also tested each method with 1 Mb long sequences (Supplementary Figure 2). As shown in Supplementary Figure 2, though every method tested exhibits higher variance when using such short sequence, eSMC exhibits a more accurate demographic history and with lower variance.

Concerning other simpler demographic scenarios (constant population size, bottleneck, expansion, decrease), eSMC shows good convergence properties although it exhibits some bias in the bottleneck scenario (Supplementary Figure 3). The bottle-neck appears smoothly estimated as a curve and not a sharp population size change. PSMC’, MSMC and MSMC2, are run with default values with the exception that the initial value of 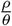 is set to 1. PSMC’ displays convergence properties similar to eSMC (Supplementary Figure 4), MSMC exhibits strong overestimation in recent time with very high effective population although a more recent demography is estimated (Supplementary Figure 5). MSMC2 shows biased results and always displays a strong overestimation in the far past (Supplementary Figure 6) and fails to detect expansions and constant demography. It is important to note that the other methods tested here all exhibit higher variances than eSMC. All estimated recombination over mutation ratios can be found in Table 2.

We now assume 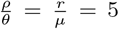, with the mutation and recombination rate respectively set to 2.5 × 10^−8^ and 1.25 × 10^−7^ per generation per nucleotide. Note that fixing 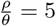 means that we estimate the demographic history conditioned on this fixed ratio (which we know is the true one). All methods provide strongly biased estimates (Supplementary Figure 7) and the estimated demographic histories are very flat. Increasing the ratio 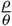 (or in this case 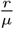) further, the estimated demographic history tends to a constant population size (results not shown). If we now allow the methods to simultaneously estimate demography and the ratio 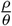, using 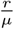 (the real value of the molecular ratio) as an input parameter (or initial value) for 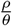, the results exhibit smaller but similar biases. However if the initial value for 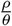 is set to 1 (or smaller) the results exhibit almost no bias (Supplementary Figure 8). In this case, the demographic history is accurately estimated although the estimated ratio 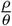 is smaller than 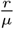 (see Table 2). While of general importance, these results have not been highlighted in previous works and we discuss their relevance later on. In the following analyses, we launch eSMC, PSMC’, MSMC and MSMC2 to estimate the demographic history and the ratio 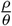 using 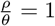 as the initial value.

#### 3.1.2 Convergence property with dormancy (seed- or egg-banks)

We first focus on sequences simulated under the “saw-tooth” scenario in the presence of seed-banks (Figure 2). Setting the mutation and recombination rates to 2.5 × 10^−8^ per generation per bp. Using eSMC, we obtain an accurate estimation of the demography (*χ*_*t*_) and of the germination rates (*β*). Under four germination rates *β* with values 1 (no seed-bank), 0.5 (two-year seed-bank), 0.2 (long-lived five-year seed-bank) and 0.1 (long-lived ten-year seed-bank), we respectively estimate an average germination rate of 0.92, 0.53, 0.22 and 0.09. As seed-banks affect the time window of the estimated demography, in their presence, inference of more ancient events is possible [59]. In models where seed-banks cannot be accounted for (PSMC’,MSMC,MSMC2), census population size is strongly overestimated when seed-banks are present. When seed-banks are long lived (*e.g. β* = 0.1, giving a mean dormancy of ten generations), eSMC slightly underestimates *β* and the population size. This is because in presence of strong seed-banks, coalescent times increase, which can lead to violation of the infinite site model. Therefore when the molecular mutation and recombination are set to 5 × 10^−9^ per generation per bp, better fits are obtained (see Supplementary Figure 9).

**Figure 2:**
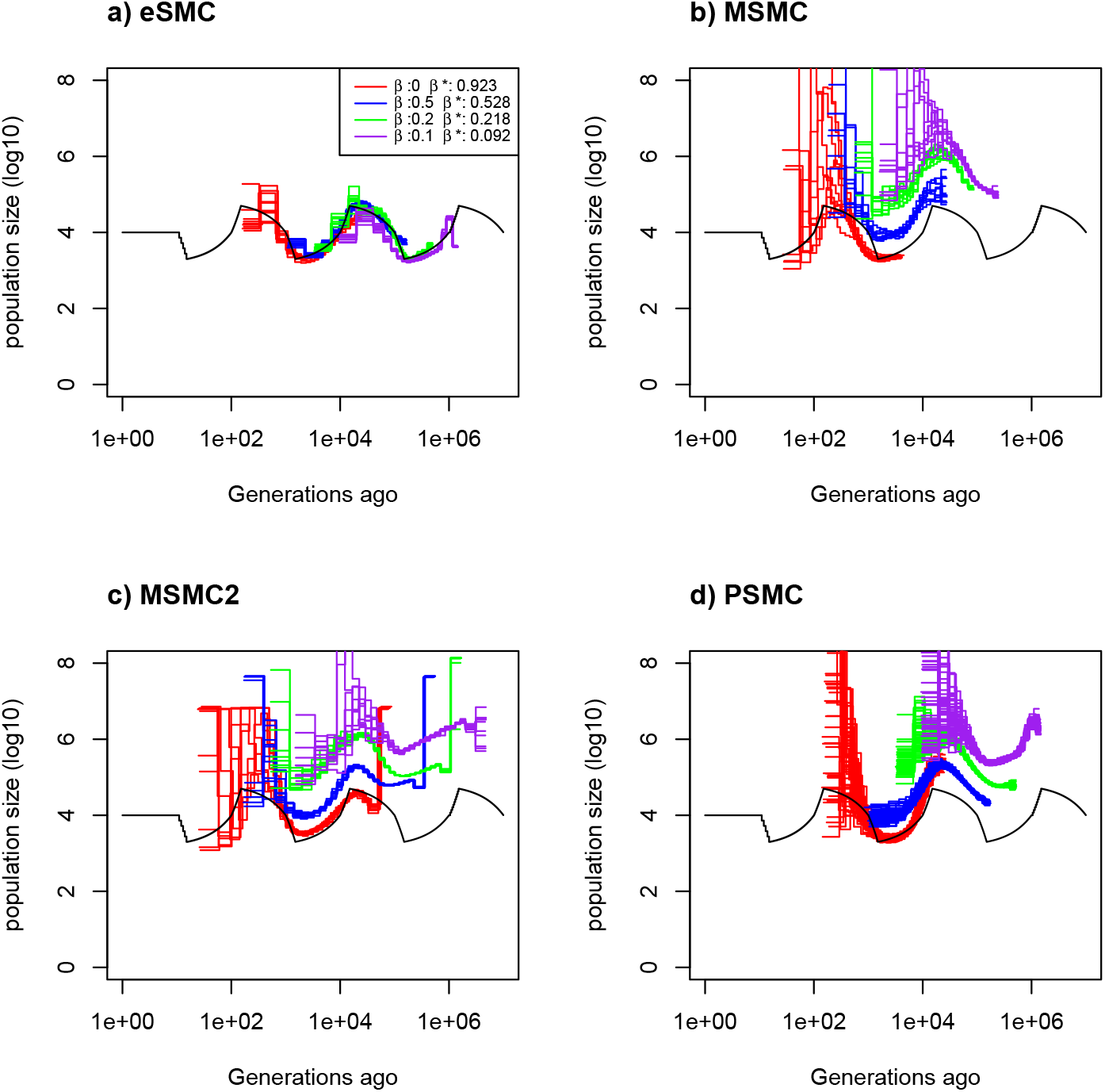
Estimated demographic history using four simulated sequences of 10 Mb and ten replicates under a saw-tooth demographic scenario (black). The mutation and recombination rates are set to 2.5 × 10^−8^ per generation per bp. Therefore 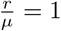. We simulate under four different germination rates *β* = 1 (red), 0.5 (blue), 0.2 (green) and 0.1 (purple), hence we respectively have 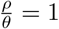, 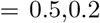 and 0.1. The demographic history is estimated using a) eSMC, b) MSMC, c) MSMC2 and d) PSMC’. *β*\* represents the estimated germination rate by eSMC.

In the simpler demographic scenarios (constant population size, bottleneck, expansion and decrease, see Supplementary Figure 10), mutation and recombination rates per site are set to 2.5×10^−8^ per generation per bp. The germination rate and the demographic histories estimated by eSMC are accurate for most of the demographic scenarios considered, except in the case of a bottleneck scenario, where the decrease in population size is not easily captured. Furthermore, in presence of strong seed-banks (*β* = 0.2 or 0.1) biases occur in the far past. Once again, this tendency disappears when the molecular mutation and recombination rates per site are lowered so as not to violate the infinite site model (*μ* and *r* = 5 × 10^−9^ per generation per bp, see Supplementary Figure 11).

#### 3.1.3 Convergence property with self-fertilization

We analyze sequences from a population simulated under the “saw-tooth” scenario with different rates of self-fertilization *σ* (Figure 3). The mutation and recombination rates are set to 2.5 × 10^−8^ per generation per bp 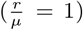. For four different self-fertilization rates *σ* = 0 (no self-fertilization), 0.5 (50% selfing), 0.8 (80% selfing)and 0.9 (90% selfing), we estimate the self-fertilization rate respectively at 0.17, 0.52, 0.78 and 0.88. In addition, we find that eSMC accurately estimates demography, while MSMC, MSMC2 and PSMC’ exhibit a small bias in the estimation of the demographic history. Neglecting self-fertilization therefore seems to be of smaller consequence than neglecting dormancy (see above), as self-fertilization has a very small impact on the the inferred demographic history. Variance in the estimations increase for higher rates of *σ*. When the mutation rate is set to 2.5 × 10^−8^ per generation per bp and the recombination rate to 1.25 × 10^−7^ per generation per nucleotide 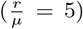, the self-fertilization rate is overestimated for small values of *σ* (Supplementary Figure 12), but well estimated for higher values of *σ*. The estimation of the demographic history remains accurate, however, though slightly biased for small values of self-fertilization. The other methods (PSMC’,MSMC,MSMC2) present stronger biases in the estimated demographic history.

**Figure 3:**
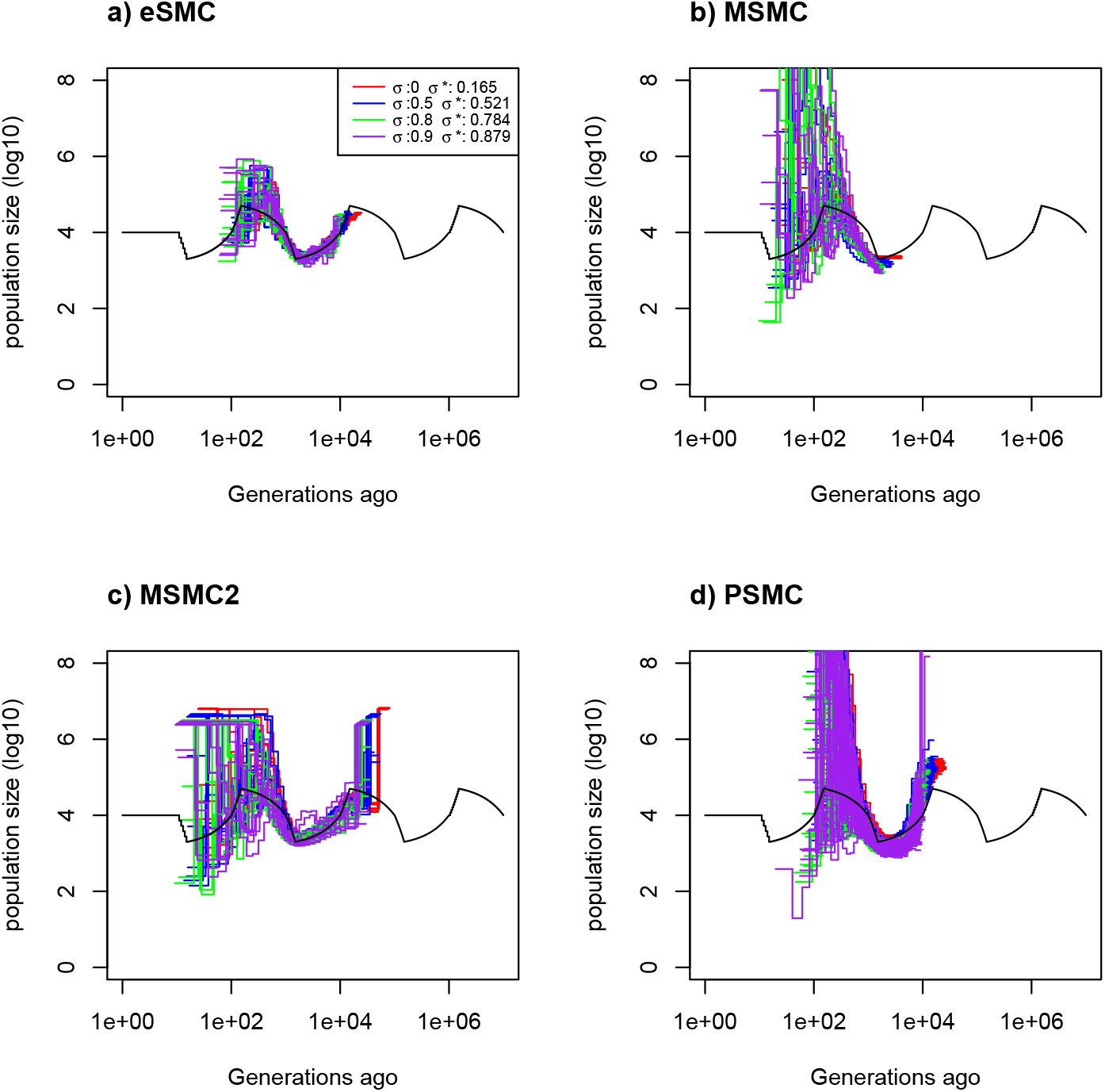
Estimated demographic history using four simulated sequences of 10 Mb and ten replicates under a saw-tooth demographic scenario (black). The mutation and recombination rates are set to 2.5 × 10^−8^ per generation per bp, and simulations were run for four different self-fertilization rates (*σ* = 0 (red), 0.5 (blue), 0.8 (green) and 0.9 (purple)), and as 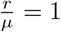, this gives 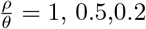 and 0.1 respectively. The demographic history is estimated using a) eSMC, b) MSMC, c) MSMC2 and d) PSMC’. *σ*\* represents the self-fertilization rate estimated by eSMC.

In the simpler demographic scenarios tested (Supplementary Figure 13), the rate of self-fertilization is estimated fairly well, though there is an impact of the considered demographic scenario. However, in absence of self-fertilization, eSMC infers a residual rate of self-fertilization (below 0.2). While, the demographic history is accurately estimated.

#### 3.1.4 Convergence property with both dormancy and self-fertilization

Self-fertilization and dormancy have opposing effects on the coalescent rate, which may decrease the inference accuracy of their respective rates when they are simultaneously present. Here we test different combinations of seed/egg-banks and self-fertilization that result in the same 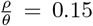 ratio where 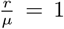 (setting mutation and recombination rates to 2.5 × 10^−8^ per generation per bp) (Figure 4). Without any prior knowledge, it is not possible for eSMC to estimate the correct set of parameters. However, by setting general “ecological” priors for either *β* or *σ* (*e.g.* 0 ≤ *β* ≤ 0.5 or 0.5 ≤ *σ* ≤ 1), eSMC can accurately infer the demographic history, though the estimations of *β* and *σ* can be slightly biased (see Figure 5). eSMC seems to overestimate *β* and *σ*, thus leading to a small overestimation of the census population size. In addition we test our model on similar data, but where recombination rate is higher (set to 1.667 × 10^−7^ per site per generation) as in our studied species the recombination rate has been estimated higher than the mutation rate [25, 39]. We thus now have 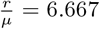 and 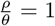. The demographic history displays smaller variance but similar estimation of self-fertilization and germination parameters (Supplementary Figure 14).

**Figure 4:**
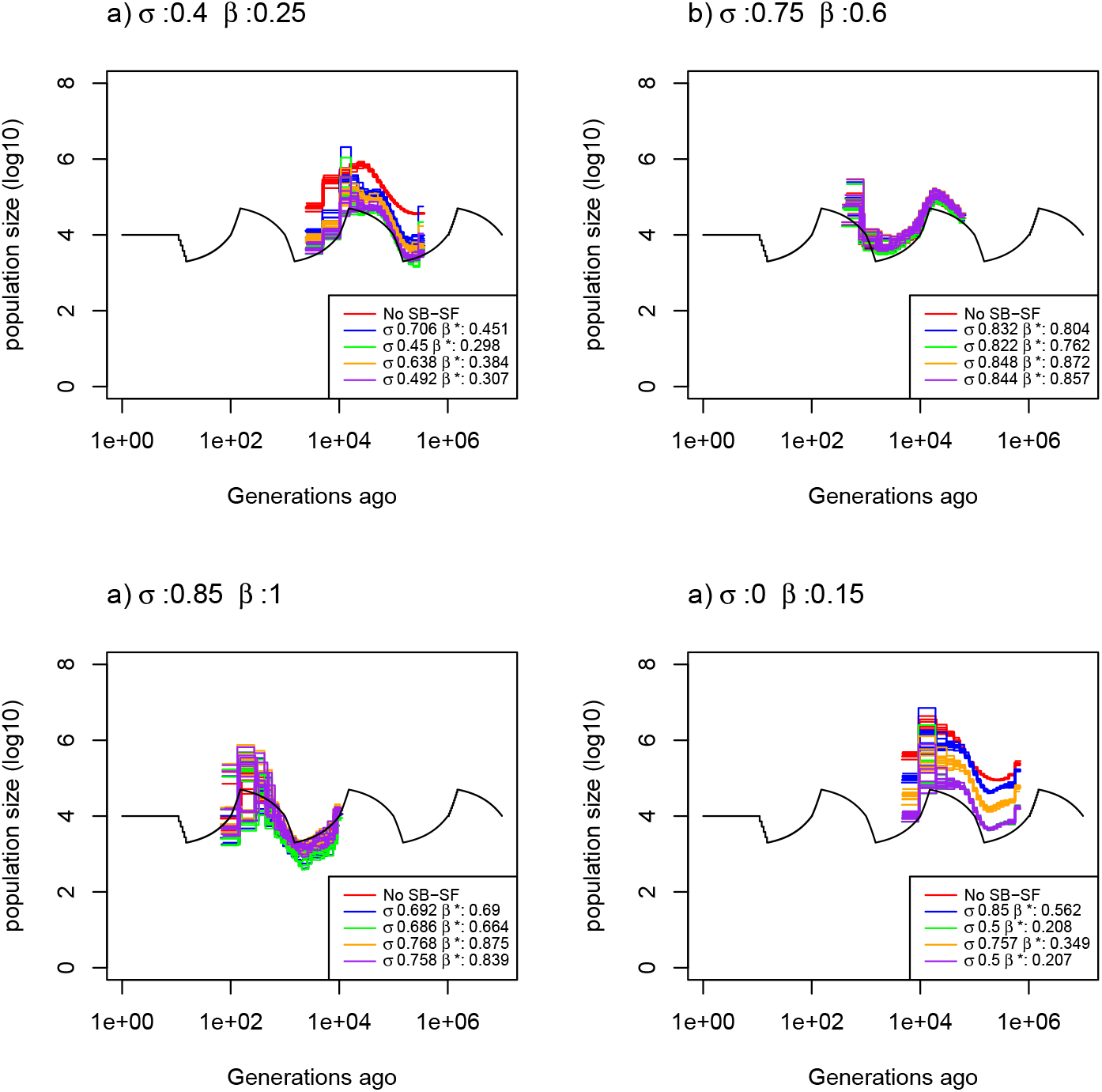
Demographic history estimated by eSMC for ten replicates using four simulated sequences of 10 Mb under a saw-tooth demographic scenario and four different combinations of germination (*β*) and self-fertilization (*σ*) rates but resulting in the same 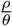. Mutation and recombination rates are set to 2.5 × 10^−8^ per generation per bp, giving 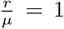. The four combinations are: a) *σ* = 0.4 and *β* = 0.25, b) *σ* = 0.75 and *β* = 0.6, c) *σ* = 0.85 and *β* = 1 and d) *σ* = 0 and *β* = 0.15. Hence, for each scenario 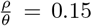 For each combination of *β* and *σ*, eSMC was launched with five different prior settings: ignoring seed-banks and self-fertilization (red), accounting for seed-banks and self-fertilization but without setting priors (blue), accounting for seed-banks and self-fertilization with a prior se3t9only for the self-fertilization rate (green), only for the germination rate (orange) or for both (purple). *σ*\* and *β*\* respectively represent the estimated self-fertilization and germination rate.

**Figure 5:**
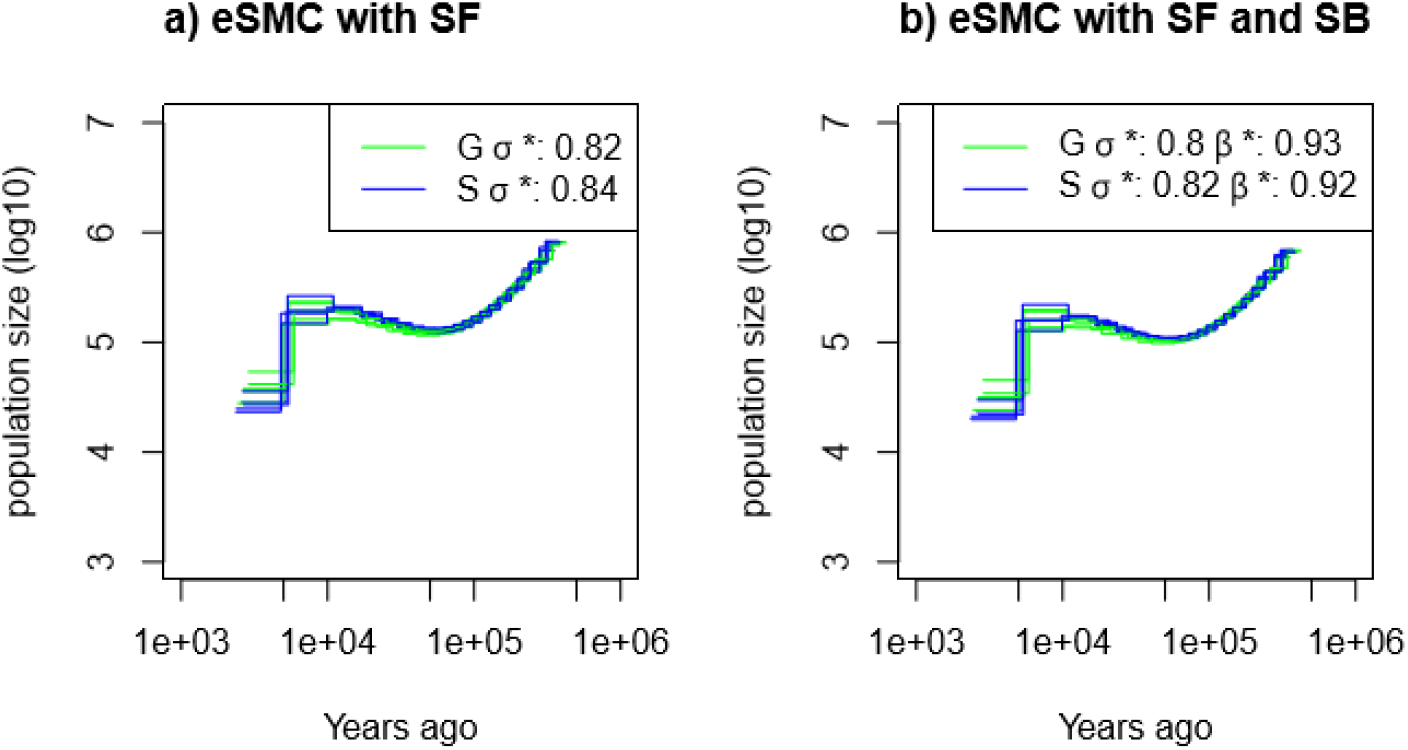
Demographic history of two European (Sweden (S, blue) and German (G, green)) populations of *A. thaliana* estimated using eSMC: a) accounting only for selfing (*σ* is a variable and *β* = 1) and b) accounting simultaneously for selfing and seed-banking (*σ* bounded between 0.5 and 0.99 and *β* bounded between 0.5 and 1). Mutation rate is set to 7 × 10^−9^ per generation per bp and recombination respectively set for chromosome 1 to 5 to 3.4 × 10^−8^, 3.6 × 10^−8^, 3.5 × 10^−8^, 3.8 × 10^−8^, 3.6 × 10^−8^) per generation per bp. *σ*\* and *β*\* respectively represent the estimated self-fertilization and germination rates.

### 3.2 Inferring self-fertilization, seed-banks and demography in *Arabidopsis thaliana*

Using 12 individual full genome sequence data obtained from two accessions of *A. thaliana* (one from Sweden and the other from Germany), we inferred the demography of each population using eSMC, PSMC’, MSMC and MSMC2 (Supplementary Figure 15). When ignoring self-fertilization, both populations have a common demographic history, similarly inferred by the different methods, except MSMC, whose results exhibit a higher variance for the Swedish population. Furthermore, we observe a non-negligible deviation between the recombination rate estimated using these inference methods 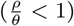 and what has been obtained using experimental approaches 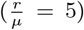 [39]. When accounting only for self-fertilization (hence imposing *β* = 1), eSMC estimates a high self-fertilization rate averaged at *σ* = 0.82 in the German population and 0.84 in the Swedish one. These rates are not as high as what has been recorded previously [44, 1]. When running analyses per chromosome, we found no significant chromosome effect on these estimations. When simultaneously estimating *β, σ* and the population size, we find a slightly lower *σ* = 0.80 in the German population and 0.82 in the Swedish one (Figure 5). Here, eSMC estimates a germination rate *β* higher than 0.9 in both populations, implying that there is no long-term seed-bank in these populations (less than 1.1 generations of dormancy on average).

### 3.3 Inferring egg-banks and demography in *Daphnia pulex*

The inferred demographic history of a single population of *D. pulex* is qualitatively similar using eSMC and PSMC’ (Figure 6). Fixing the self-fertilization rate at *σ* = 0, the inferred mean generation time before the hatching of dormant eggs produced by sexual reproduction depends on the number of parthenogenetic cycles that occur on average per year. Here one generation is considered to be of one cycle of asexual or sexual reproduction, hence several generations can occur in a single year. For this specific population, a maximum of five parthenogenetic cycles before sexual reproduction is assumed [25]. It is important to note that the number of parthenogenetic cycles can affect the ratio 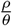. Therefore we tested the effect of the value of the average number of parthenogenetic cycles on the estimation of the germination rate. Even for the extreme cases (no parthenogenesis and five parthenogenetic cycles), eSMC detects dormancy, with *β* < 0.3. Hence we can bound the average dormancy to be between 3 and 18 generations, revealing the existence of at least moderate dormancy in this species.

**Figure 6:**
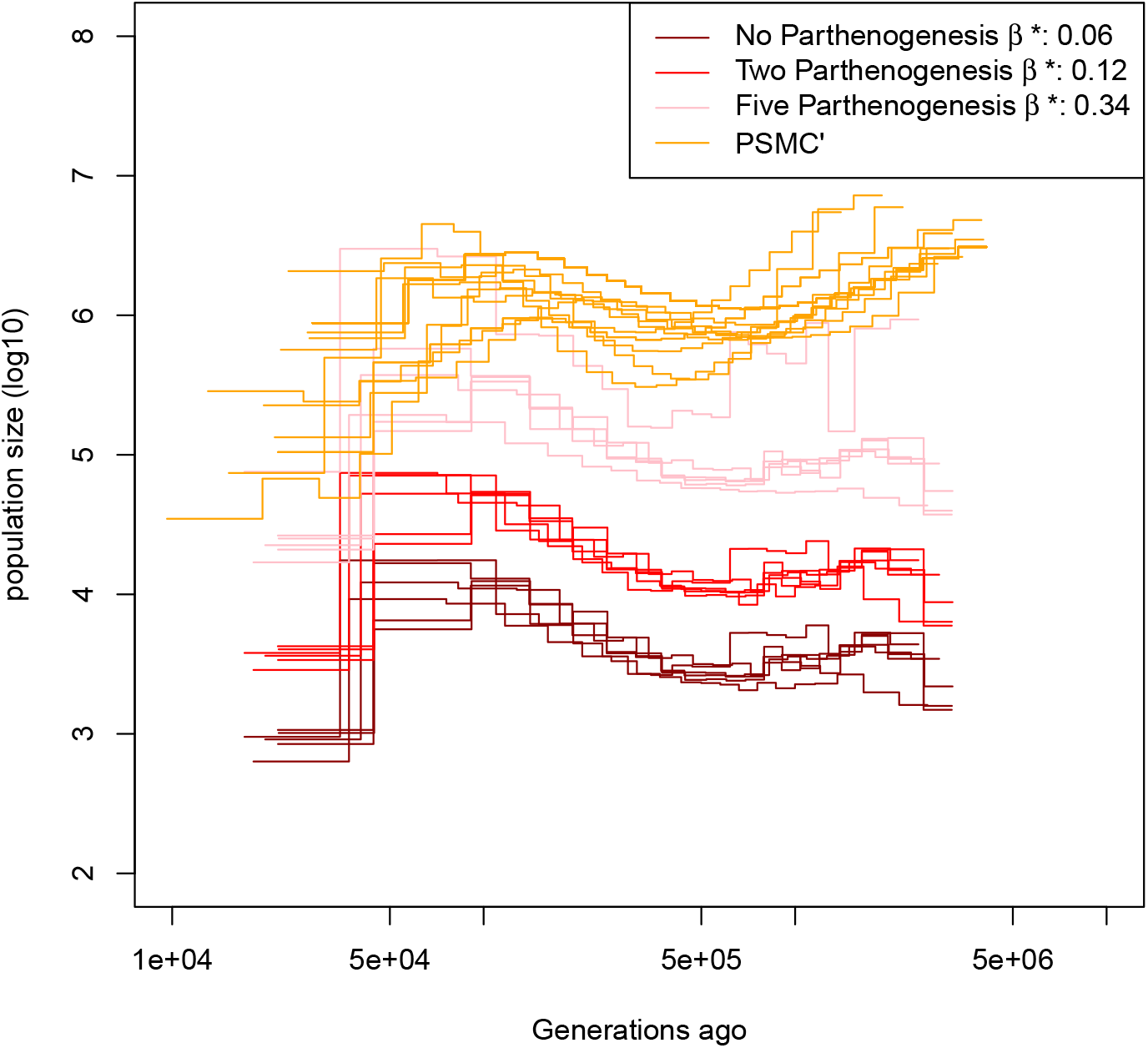
Demographic history estimated by eSMC on six individuals of *D. pulex* accounting for egg-banks (*β* is a variable and *σ* = 0). Different assumptions concerning the number of parthenogenetic cycles before the production of the dormant egg are made: Five cycles (pink), two cycles (red) and no parthenogenesis (dark red). Mutation and recombination rates are respectively set to 4.33×10^−9^ and 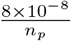 per generation per bp, where *n*_*p*_ is the number of reproductive cycles per year, parthenogenetic and sexual.

## 4 Discussion

The existing statistical inference methods based on full genome polymorphism data estimate the past demographic history under assumptions of the model which can be violated by many species. Here, we develop a method where ecological and life history traits can not only be included, but can also be inferred from sequence data along with the past demography. We demonstrate the capacity of our method to accurately recover the germination rate (and therefore the presence and strength of dormancy). Ecology and life history traits can affect *ρ* and *θ* differently which can be detected by our HMM through the estimation of 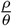. Hence, one requires knowledge on the molecular ratio of recombination over mutation 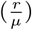. Furthermore, simulated results tend to show that violation of the infinite site assumption leads to a small underestimation of the germination rate. In a similar way, we show that our model can also retrieve the self-fertilization rate. In simulated results, we find that higher self-fertilization rate (*e.g.* 0.9) increases the variance of the estimation. Hence more data are required to increase accuracy. In its current version, our model cannot disentangle the genomic signatures of self-fertilization and seed-banks, and tends to favor self-fertilization over seed-banks. However, this bias can be reasonably avoided if some prior knowledge about the species biology and ecology is available. For example in the case of seed-banks (or egg-banks), many species with long lived dormant stages (*e.g.* defined as longer than five years in plants) have been identified based on field data and are found in reference databases [5, 6]. For self-fertilization, many species exhibiting self-fertilization are also reported, or at least the type of reproduction mode (selfing, partial outcrossing, outcrossing). Setting priors (*β* and/or *σ* < or > 0.5) can thus reflect this common *a priori* knowledge.

We demonstrate that, even in the absence of self-fertilization and dormancy, the accuracy of our model (eSMC) in inferring demographic history is similar to, and in several cases better than, other methods that are widely used. This is because the model runs all pairwise analysis, increasing the amount of available information and the capacity to bound estimated values. It is important to note that abrupt or sudden scenarios, such as bottlenecks, are difficult to infer, irrespective of the method used [35]. The effect of sharp (sudden) demographic events on allele frequencies can in fact be delayed for small sample size, explaining why the demographic history of the bottleneck is smoothed as hinted in [52]. Because our model can efficiently integrate information from several sequences and chromosomes, it gives good convergence properties with only a few sequences, that do not need to be very long (a total of 10 Mb is sufficient, yet each scaffold should be longer than 1 Mb). eSMC therefore allows an accurate estimation of demographic history with just one or two sequenced individuals and when using this method the quality of the genetic sequences should prevail over their quantity (in this our method is similar to the PSMC’).

Throughout the paper we have highlighted two main ratios that are of great importance when using inference methods: the ratios 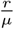 and 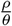, respectively the per site molecular and effective ratios of the recombination over the mutation rate. We have used the deviation between 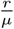 and 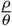 to estimate the self-fertilization or germination rate. We also show that the demographic history can contribute to a departure of 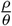 from 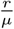. Indeed, care must be taken concerning the initial value used for 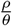. If the initial value set for 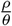 is greater than one, the inferred demographic history will be flattened, regardless of the actual value of 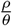. Furthermore, if the true value of 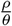 is indeed greater than one, similar biases are expected, such as flattened demographic history. The reason for this result, which applies to all species and irrespective to whether they have seed-banks or present self-fertilization, is that, if the ratio 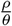 is high, the power of estimations becomes low as too few SNPs are present between the recombination spots on the genome. Therefore, not enough information is available to correctly reconstruct the local coalescent trees. We highlight that the importance of 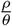 for inference was hinted in [48], but has been ignored in the literature on SMC-based methods though this ratio significantly alters the accuracy of inference when it is greater than one.

When applying eSMC to sequences data of *A. thaliana*, we find evidence of strong self-fertilization with an estimated selfing rate around 0.8. However, this rate is slighlty smaller than what is known empirically for this species, where the current rate of self-fertilization has been estimated at 0.99 [1]. Her, we speculate about three scenarios explaining this deviation. First, one possible explanation is that *A. thaliana* has most probably evolved from outcrossing to highly self-fertilizing less than 400 thousand years ago [44]. As our demographic inference dates further in the past, self-fertilization would have most likely appeared within the time window of the inferred demographic history. As a consequence, our estimate of self-fertilization (constant in time) reflects the average effect of the varying real selfing rate within the time window. Second, the under-estimation may be due to limits of the self-fertilization model, which accounts only for homologous recombination events. Yet, other types of recombination or chromosomic re-arrangements do occur in genomes. These non-accounted for mechanisms could increase the signature of recombination leading to underestimation of the self-fertilization rate. Third, we infer the self-fertilization rate of a single population in isolation (Germany or Sweden) while the past demography of *A. thaliana* consists of episodes of admixture, migration and recolonization from glacial refugia [7, 50], all of which are ignored in our model. The resulting complex population structure likely has an effect on our estimates (see discussion in [38]).

It has long been observed and/or suspected that many *Daphnia* species, in particular *D. pulex*, have resting egg-banks. The sequences analyzed using eSMC agree with this hypothesis, as we found strong evidence of dormancy. The inferred duration of dormancy greatly depends on the number of parthenogenetic generations between sexual reproduction events. Indeed, parthenogenetic cycles increase the number of mutations compared to recombination events. If we take two extreme scenarios for the specific sampled population (no parthenogenesis versus 5 generations of parthenogenesis [25]) we find a duration of dormancy between 3 and 18 generations. This is in agreement with empirical observations [10, 6], and confirms the major role of egg-banks in maintaining diversity in this species. The sequences used here originate from an intermittent standing water body (*i.e.* non-permanent pond), and populations in such environments are expected to have both higher rates of sexual reproduction as well as longer egg-banks [10]. It would therefore be interesting to test the existence of egg-banks and assess the germination rates in several *Daphnia* species and from different standing water bodies (permanent to intermittent). Our method therefore presents a way forward to the detection of egg/seed/spore-banks of many invertebrates, plant and fungal species, as well as their past demographic history using sequence data (as experimental validation of dormancy is difficult to obtain [45, 46]).

Throughout this work, we have highlighted the importance of 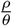 when analyzing whole genome sequence data, as it is often confused with, or considered equivalent to, 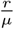. Indeed, its impact as a variable/parameter is often ignored when estimating demographic history, despite the biases it can engender when it is too large or if the wrong priors are used. Our method represents a first step for the integration of ecological traits in whole genome sequence analysis through this ratio 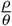. We nevertheless advise caution when using our proposed, or other HMM methods, for the inference of demography, as some assumptions may still be violated. For example we assume that mutations occur in the seed/egg-bank (a consequence of DNA damage) at the same rate as in the active population. While there is support in plants for this hypothesis [53, 8], we have no knowledge of supporting data in *Daphnia*. If mutations occur in seeds/eggs at a slower rate, we predict the estimation of dormancy to be a conservative lower bound, meaning that the seed/egg-bank is actually longer lived (*e.g.* [45]). Furthermore, additional mechanisms and processes can influence 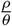 in such a way that it can be interpreted as dormancy or self-fertilization. It is therefore highly recommended that some knowledge on the biology of the species be collected before launching eSMC, or other methods, so as to define the appropriate priors.

In conclusion, the presented method is the first, to our knowledge, allowing the joint estimation of life-history traits and past demographic history based on full genome data. It is specifically adapted to the many species presenting violations of the classic Wright-Fisher model, and can be used to study the evolution of seed/egg-banking as an important bet-hedging strategy with large consequences on the rate of genome evolution [46].

## 5 Material and Method

### 5.1 Method

#### 5.1.1 ecological Sequentially Markovian Coalescent (eSMC)

The eSMC is a Hidden Markov Model along two phased haplotypes. It is an extension to the PSMC’ algorithm [40]. It adds the possibility of taking seed-banks and self-fertilization into account and simultaneously estimating their rates along with the demographic history. As in PSMC’, we assume neutrality, an infinite site model and a piece-wise constant population size. To define our HMM we need to precisely define all the following objects: the signal (observed data), the hidden states (coalescent time), the emission probabilities (probabilities of observing the data conditional to the hidden states), transition probabilities (probabilities of jump from one hidden state to another) and the probabilities of the initial hidden states. The demonstrations of the results presented here can be found in the Appendix section 1.

The signal (or observed data) depends on the hidden state and is a chain of 0s and 1s. To construct this signal, as in PSMC’, two genome sequences are compared; if, at a given position, the two nucleotides are the same on both sequences, this is indicated by a 0, otherwise by a 1 (Figure 1). As is necessary in HMM, the number of hidden states (or the coalescent times) must be finite, which is achieved by discretizing time. Therefore, the hidden state at one position is *α* if the coalescent time between the two phased nucleotides at that position is between *T*_*α*_ and *T*_*α*+1_. Given the model parameters, we know the expected coalescent time (which is 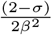), and we define *T*_*α*_:

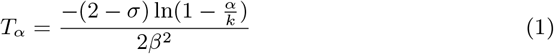

Here, *k* is the number of hidden states and *α* is an integer value between 0 and *k* − 1. *σ* and *β* are respectively the self-fertilization and the germination rate.

The emission probability *P* is the probability of observing the signal (chain of 0’s and 1’s) conditional to the hidden states (coalescent time). As in the PSMC’ algorithm, we consider an infinite site model. The emission rate is therefore given by:

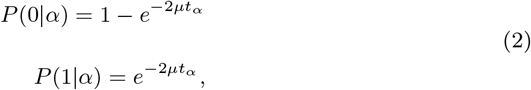

Where *μ* is the mutation rate per base pair and *T*_*α*_ the expected coalescent time in interval *α*. We find:

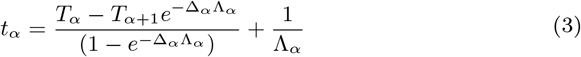

With:

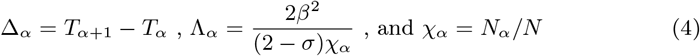

Where Δ_*α*_ is the duration (in coalescent time) of interval *α*, Λ_*α*_ is the coalescent rate in the time window *α*, *N* is the population size at time 0 and *N*_*α*_ is the population size during the time interval *α*. Using *N* and *N*_*α*_, we can calculate *χ*_*α*_ which represents the variation of population size over time. It is this value that is inferred by the model.

The transition probabilities are the probabilities of going from one hidden state to another. We find:

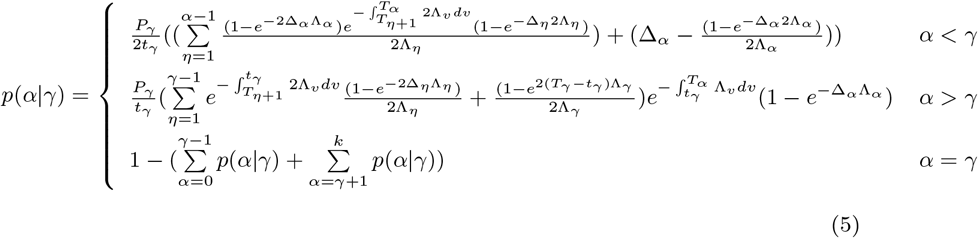

Where *P*_*γ*_ is the recombination probability between two base-pairs:

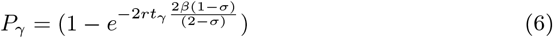

The initial probability corresponds to the first state probability. We assume this probability to be the equilibrium probability *q*_*o*_(*α*) (probability of being in state *α* at the first position). We find:

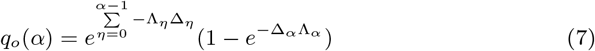

### 5.2 Material

#### 5.2.1 Simulated (pseudo-observed) Sequence data

Throughout this paper we use five different demographic scenarios: 1) constant population size, 2) expansion, 3) bottleneck and recovery, 4) decrease and 5) “saw-tooth” (a succession of expansions and decreases). These scenarios are simulated for different combinations of the self-fertilization rate (0 ≤ *σ* ≤ 0.9) and the germination rate (0.1 ≤ *β* ≤ 1). Different sequence lengths are tested, as are combinations of mutation and recombination rates. To simulate our data, we use a modified version of the coalescent simulation program scrm [42]. This modified version integrates seed-banking (or egg-banking) and self-fertilization. The simulator is available on our GitHub repository (https://github.com/TPPSellinger/escrm). All the command lines can be found in Table 1 of the Appendix. On all the simulated data, four different algorithms are used to estimate demographic history and recombination rate: our algorithm eSMC, which we compare to PSMC’, MSMC and MSMC2. PSMC, MSMC and MSMC2 are run with default parameter and 1 as initial value for 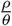.

#### 5.2.2 Sequence Data

We use 12 whole genome sequences (hence all five chromosomes) of European *A. thaliana* from the 1001 genome project [7, 50], six individuals sampled in Sweden (id: 5830, 5836, 5865, 6077, 6085 and 6087) and six from Germany (id: 7231, 7250, 7255, 7337, 7415 and 7419). Each individual is considered haploid because of very high levels of homozygosity [15]. We obtained polymorphism data (that is processed vcf files) from the authors of the study [15]. The mapping to the reference genome and SNP call was performed based on the pipeline in [15]. The mutation rate is set at 7 × 10^−9^ per generation per bp [34] and the chromosome specific recombination rates are 3.4 ×10^−8^, 3.6 × 10^−8^, 3.5 × 10^−8^, 3.8 × 10^−8^, 3.6 × 10^−8^ per generation per bp for chromosome 1 to 5 respectively [39]. We first run the four different algorithms to estimate the demographic history and recombination rate (ignoring self-fertilization and seed-banks for eSMC). Analysis are run per chromosome (represented by the different lines in the figures). We then analyse the data again with eSMC, first accounting only for self-fertilization (*β* is fixed to 1 and *σ* is estimated), and then accounting for both self-fertilization and seed-banks using reasonable priors (0.5 ≤ *β* ≤ 1 and 0.5 ≤ *σ* ≤ 1).

To infer the demographic history and the dormancy rates of *D. pulex*, we use six whole genome sequences from [25](id: SRR5004865, SRR5004866, SRR5004867, SRR5004868, SRR5004869 and SRR5004872) which are available under the accession SAMN06005639 in the NCBI Sequence Read Archive (SRA). We used the reference genome assembly PA42 v3.0 which is available at the European Molecular Biology Laboratory (EMBL) nucleotide sequencing database under accession PRJEB14656 [57]. The raw data was first trimmed using bbtools to remove duplicates, trim adapters, remove synthetic artifacts, spike-ins and perform quality-trimming based on minimum read quality of 40. Then we mapped reads using bwa (default parameters) onto the reference genome [21]. We used Samtools to convert sam to bam files [23] and GATK to remove PCR duplicates and perform local realignment around indels [29]. We used freebayes to call the SNPs and vcftools for post-processing (filtering). The pipeline is available upon request. Note that the reference genome consists of 1,822 scaffolds (average length of 85,849) and thus to avoid bias in the analyses, we only kept scaffolds above 1 Mb retaining only the 19 largest scaffolds. As the phasing quality could not be guaranteed, we only analyze the sequence data of each individual separately *D. pulex*. Indeed if sequences are phased 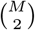 analysis can be carried out with M diploid individuals. However, if data are not phased, M analysis need to be carried out, which are the M diploid individuals. The mutation rate is set at 4.33 × 10^−9^ per generation per bp [14] and the recombination rate at 8 × 10^−8^ per event of sexual reproduction per bp [56, 25]. So as to account for the number of generations before sexual reproduction takes place, the recombination rate is re-scaled by *n*_*p*_ which represents the total number of generations per year. If we consider *n*_*p*_ = 5, the recombination rate is scaled by 0.2 [25]. We also test how the number of parthenogenetic generations between sexual reproductive events could affect the quality of the inference, and rescale the recombination rate accordingly. The scenarios we test are: no parthenogenesis, two generations of parthenogenesis and five generations of parthenogenesis, therefore rescaling the recombination rate by 1, 0.5 and 0.2 respectively. The sequences of each individual are analyzed with PSMC’ and eSMC only, as MSMC and MSMC2 require accurate and reliable phasing, which is not the case for these sequences. We then account for egg-banks using eSMC and imposing no priors on *β* and setting *σ* = 0. The multihetsep files for *A. thaliana* and *D. pulex* analysis are available on the GitHub repository (https://github.com/TPPSellinger/Daphnia_pulex_data and https://github.com/TPPSellinger/Arabidopsis_thaliana_data).

## Supporting information

Appendix Mathematical derivations

