## Supplementary Figures for "Inference of past demography, dormancy and self-fertilization rates from whole genome sequence data"

### Appendix

Thibaut Sellinger[1], Diala Abu Awad[1], Markus Möst[2], Aurélien Tellier[1]
[1] Department of Population Genetics, Technische Universität München
[2] Department of Ecology, University of Innsbruck

July 12, 2019

#### 1 Supplementary Figures

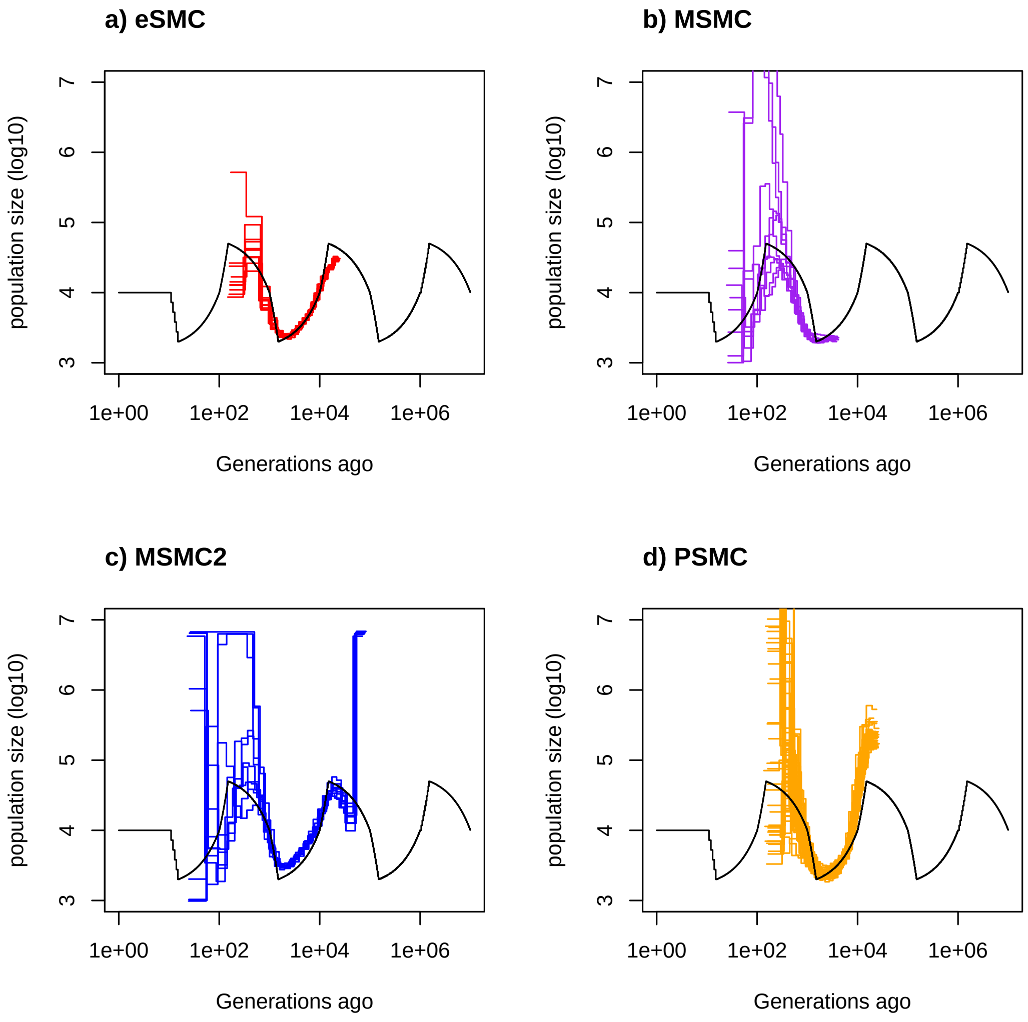


Figure 1: Estimated demographic history using four simulated sequences of 10Mb under a saw-tooth demographic scenario with 10 replicates. Mutation and recombination rate are set to $2.5 \times 10^{-8}$ per generation per bp. Therefore $\frac{r}{\mu} = \frac{\rho}{\theta} = 1$. The simulated demographic history is represented in black. a) Demographic history estimated by eSMC (red). b) Demographic history estimated by MSMC (purple). c) Demographic history estimated by MSMC2 (blue). d) Demographic history estimated by PSMC' (orange).

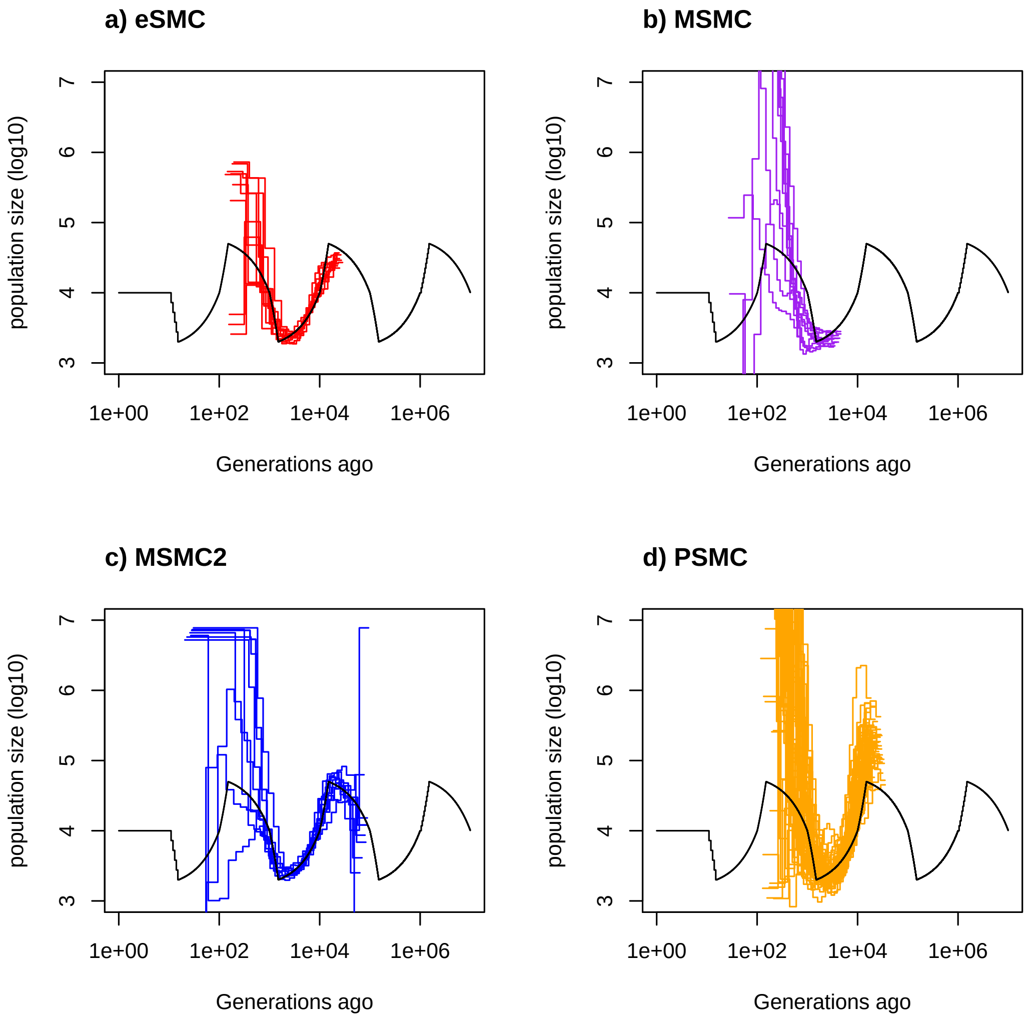


Figure 2: Estimated demographic history using four simulated sequences of 1Mb under a saw-tooth scenario with 10 replicates. Mutation and recombination rate are set to $2.5 \times 10^{-8}$ per generation per bp. Therefore $\frac{r}{\mu} = \frac{\rho}{\theta} = 1$. The simulated demographic history is represented in black. a) Demographic history estimated by eSMC (red). b) Demographic history estimated by MSMC (purple). c) Demographic history estimated by MSMC2 (blue). d) Demographic history estimated by PSMC' (orange).

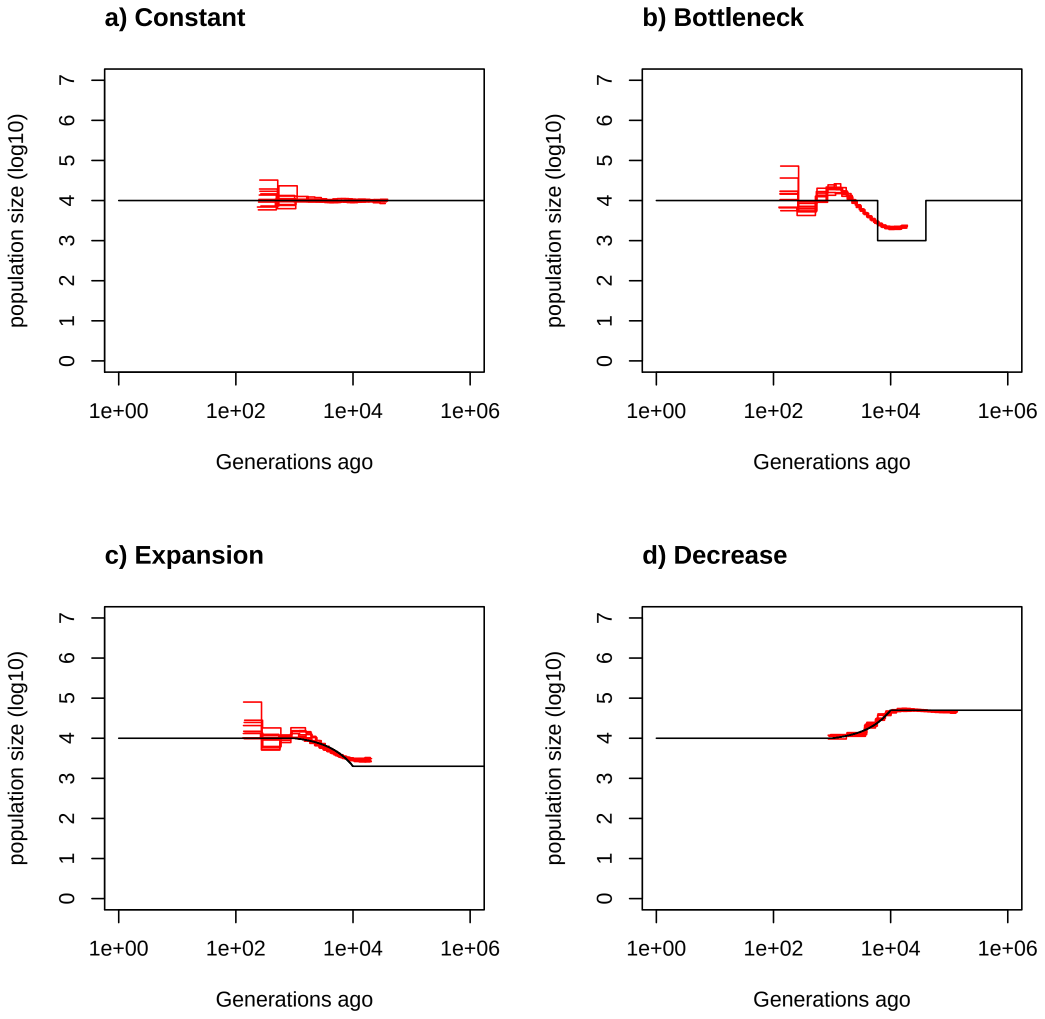


Figure 3: Estimated demographic history using four simulated sequences of 10Mb under 4 different demographic scenarios with 10 replicates. Mutation and recombination rate are set to $2.5 \times 10^{-8}$ per generation per bp. Therefore $\frac{r}{\mu} = \frac{\rho}{\theta} = 1$. The simulated demographic history is represented in black. a) Demographic history simulated under a constant population size. b) Demographic history simulated under a bottleneck. c) Demographic history simulated under an expansion. d) Demographic history simulated under a decrease. Demographic history estimated by eSMC is in red.

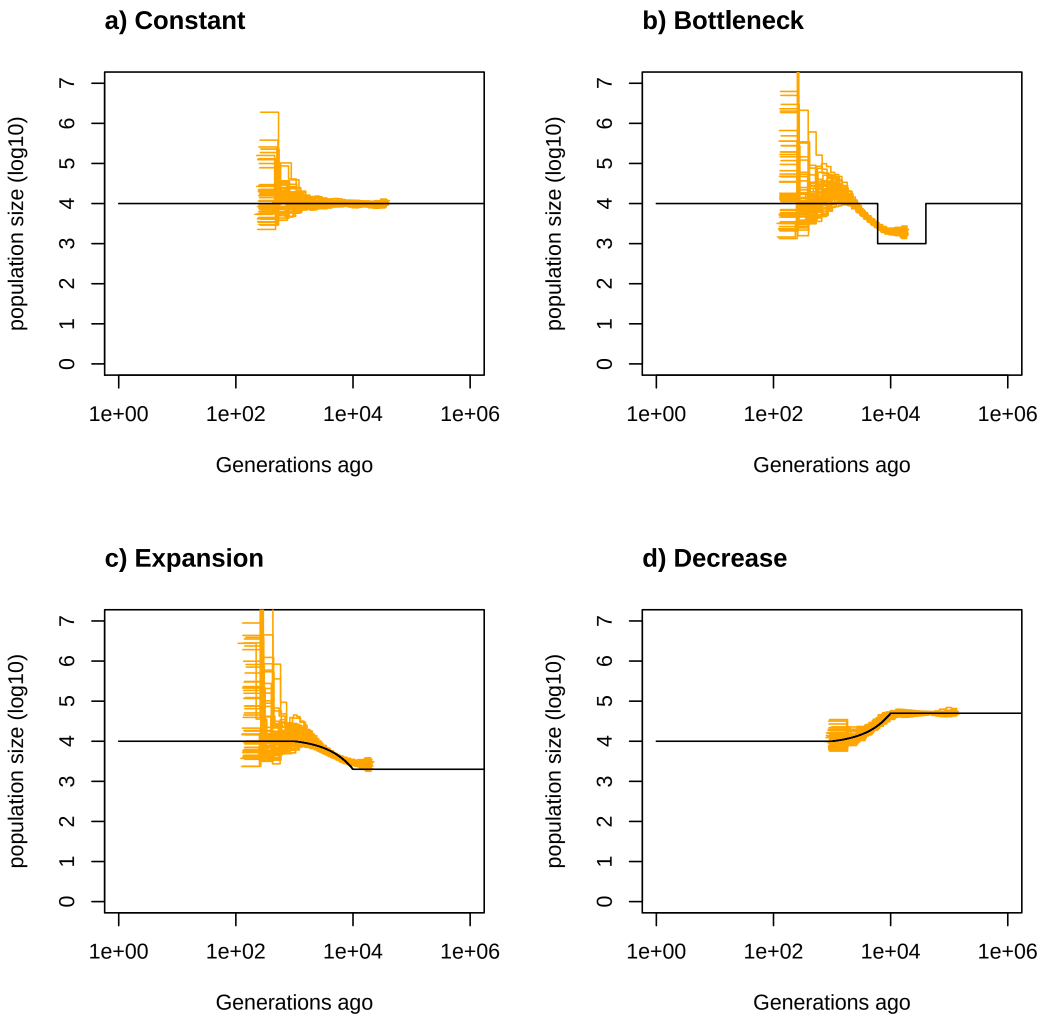


Figure 4: Estimated demographic history using four simulated sequences of 10Mb under 4 different demographic scenarios with 10 replicates. Mutation and recombination rate are set to $2.5 \times 10^{-8}$ per generation per bp. Therefore $\frac{r}{\mu} = \frac{\rho}{\theta} = 1$. The simulated demographic history is represented in black. a) Demographic history simulated under a constant population size. b) Demographic history simulated under a bottleneck. c) Demographic history simulated under an expansion. d) Demographic history simulated under a decrease. Demographic history estimated by PSMC' is in orange.

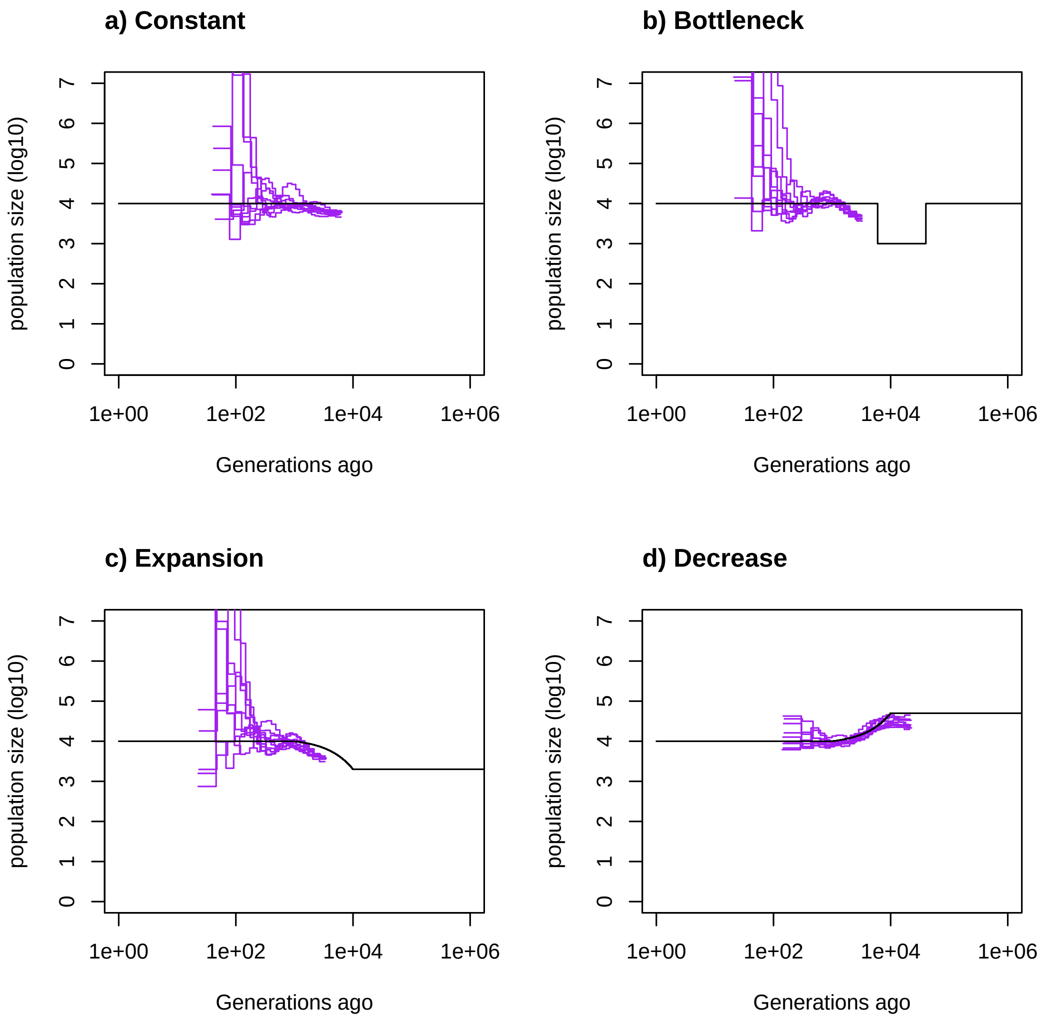


Figure 5: Estimated demographic history using four simulated sequences of 10Mb under 4 different demographic scenarios with 10 replicates. Mutation and recombination rate are set to $2.5 \times 10^{-8}$ per generation per bp. Therefore $\frac{r}{\mu} = \frac{\rho}{\theta} = 1$. The simulated demographic history is represented in black. a) Demographic history simulated under a constant population size. b) Demographic history simulated under a bottleneck. c) Demographic history simulated under an expansion. d) Demographic history simulated under a decrease. Demographic history estimated by MSMC is in purple.

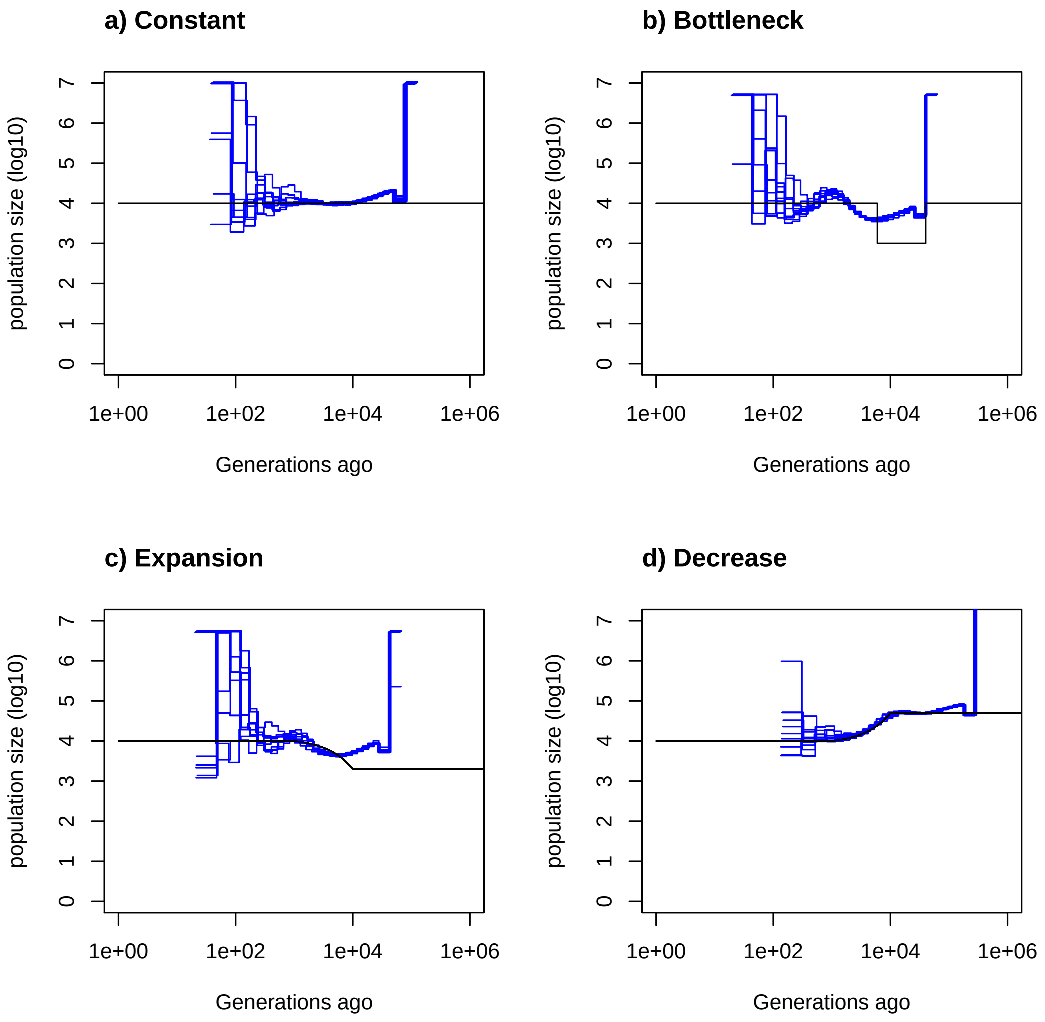


Figure 6: Estimated demographic history using four simulated sequences of 10Mb under 4 different demographic scenarios with 10 replicates. Mutation and recombination rate are set to $2.5 \times 10^{-8}$ per generation per bp. Therefore $\frac{r}{\mu} = \frac{\rho}{\theta} = 1$. The simulated demographic history is represented in black. a) Demographic history simulated under a constant population size. b) Demographic history simulated under a bottleneck. c) Demographic history simulated under an expansion. d) Demographic history simulated under a decrease. Demographic history estimated by MSMC2 is in blue.

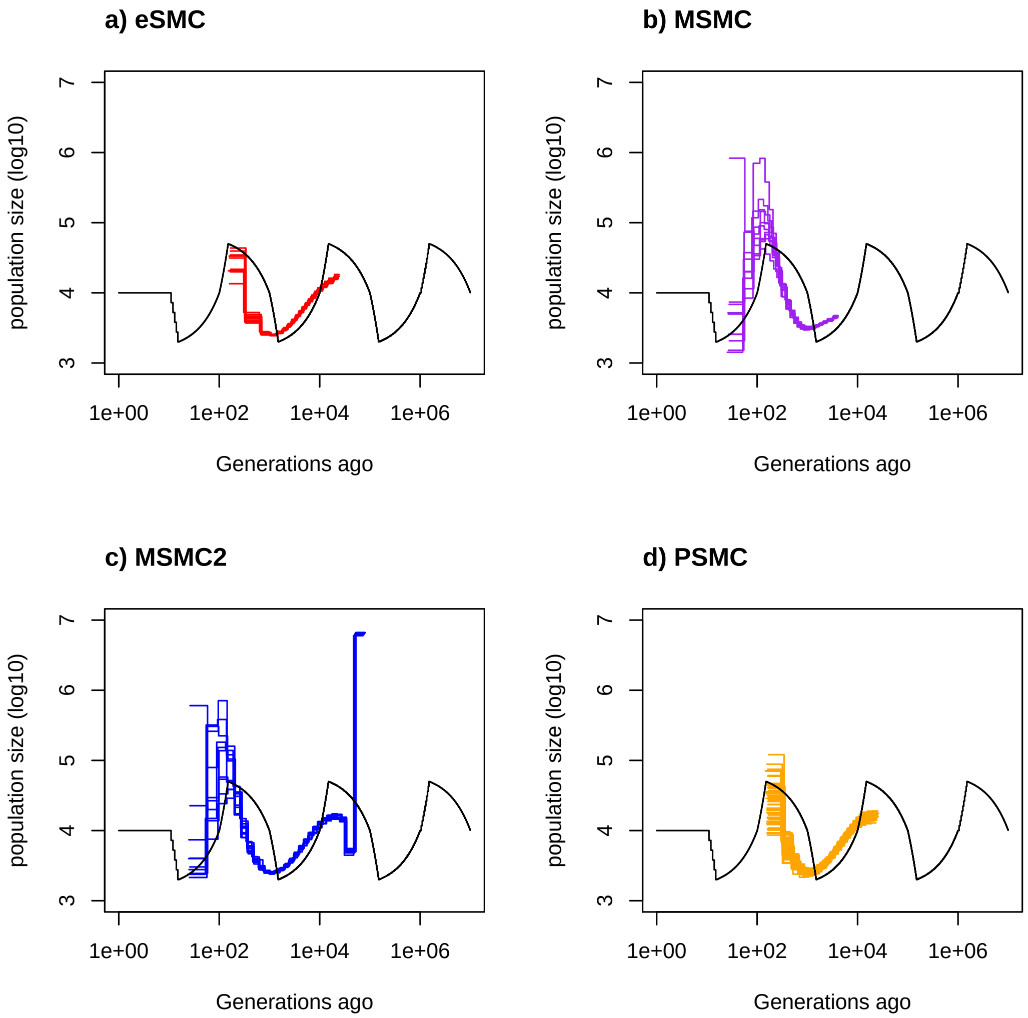


Figure 7: Results are obtained by fixing recombination rate to real value. Estimated demographic history using four simulated sequences of 10Mb under a saw-tooth scenario with 10 replicates. Mutation and recombination rate are set to $2.5 \times 10^{-8}$ and $1.25 \times 10^{-7}$ per generation per bp. Therefore $\frac{r}{\mu} = \frac{\rho}{\theta} = 5$. The simulated demographic history is represented in black. a) Demographic history estimated by eSMC (red). b) Demographic history estimated by MSMC (purple). c) Demographic history estimated by MSMC2 (blue). d) Demographic history estimated by PSMC' (orange).

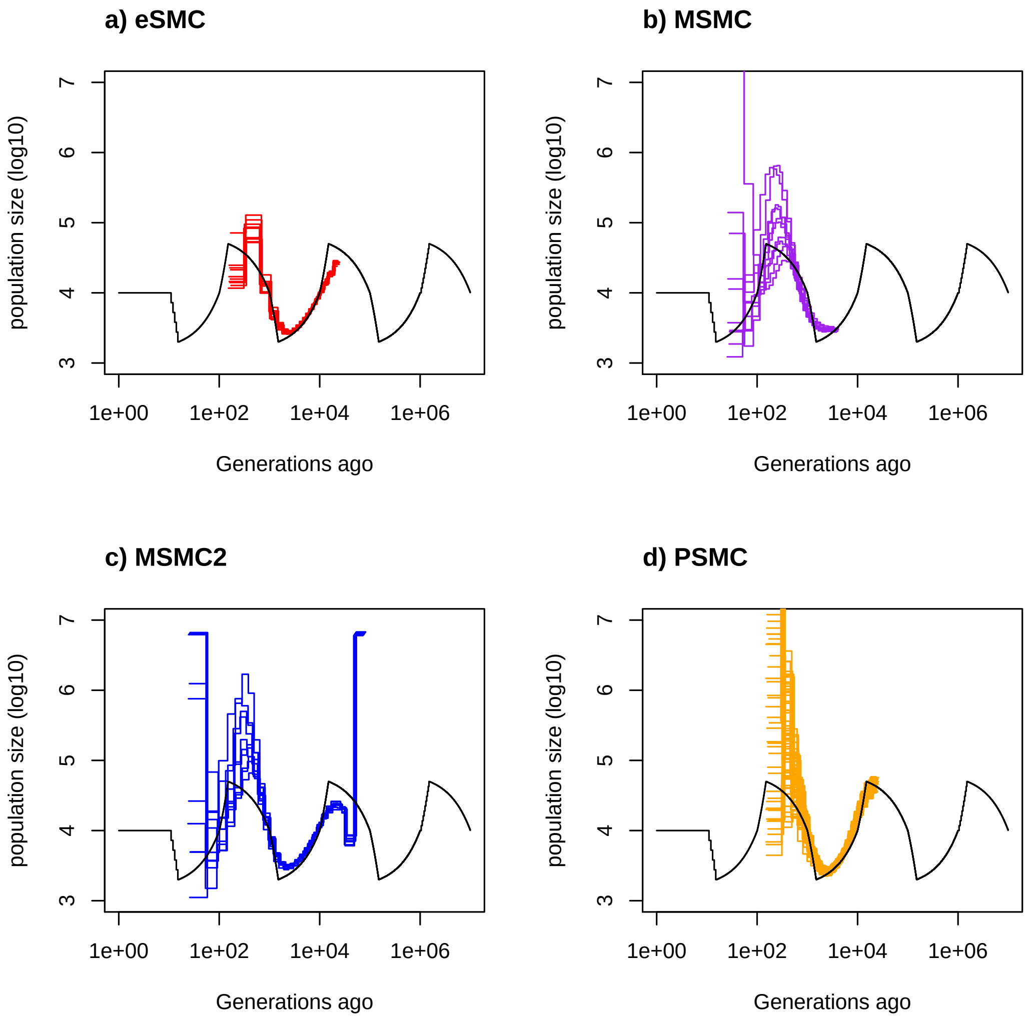


Figure 8: Results are obtained by estimating recombination rate with initial value equal to mutation rate ($\frac{\rho}{\theta} = 1$). Estimated demographic history using four simulated sequences of 10Mb under a saw-tooth scenario with 10 replicates. Mutation and recombination rate are set to $2.5 \times 10^{-8}$ and $1.25 \times 10^{-7}$ per generation per bp. Therefore $\frac{r}{\mu} = \frac{\rho}{\theta} = 5$. The simulated demographic history is represented in black. a) Demographic history estimated by eSMC (red). b) Demographic history estimated by MSMC (purple). c) Demographic history estimated by MSMC2 (blue). d) Demographic history estimated by PSMC' (orange).

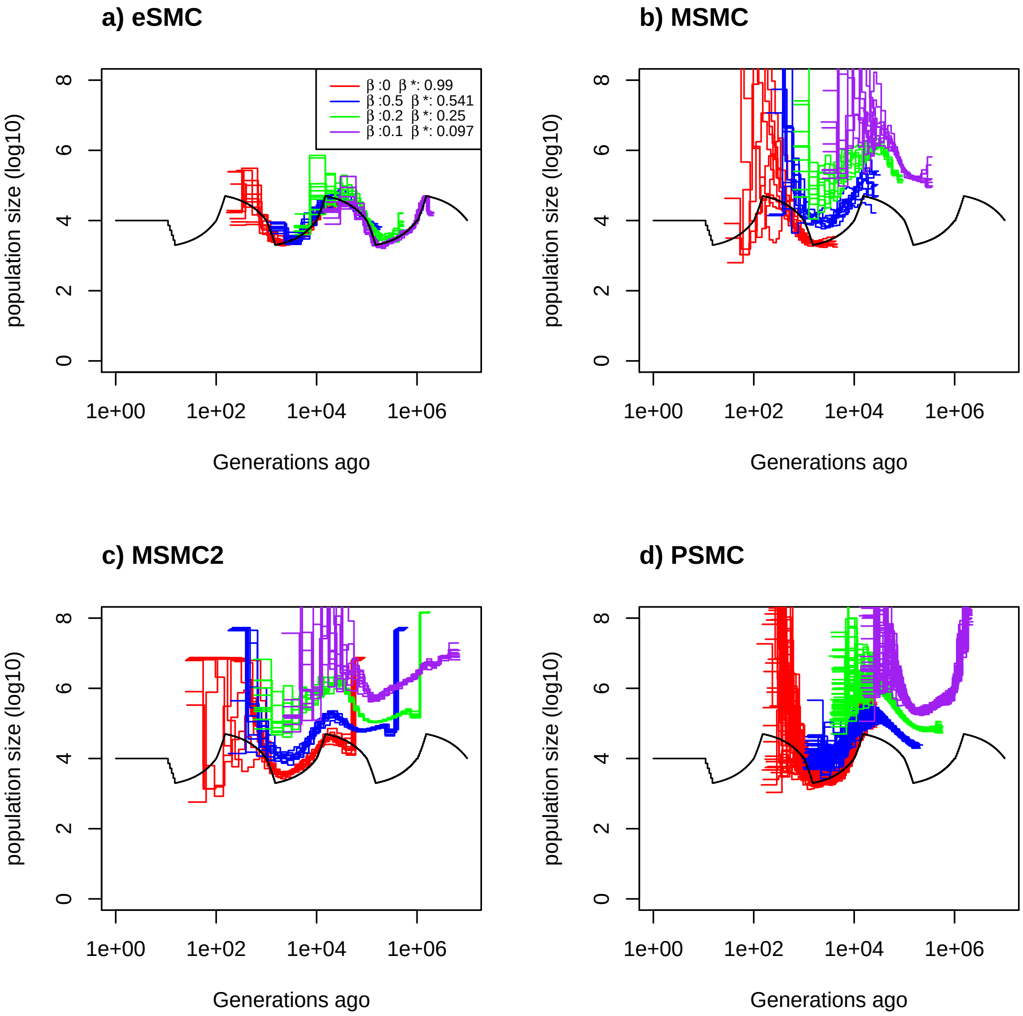


Figure 9: Estimated demographic history using four simulated sequences of 10Mb and ten replicates under a saw-tooth demographic scenario (black). Simulation were done under four different germination rate $\beta$ (1,0.5,0.2 and 0.1). The mutation and recombination rates are set to $5 \times 10^{-9}$ per generation per bp. Therefore $\frac{r}{\mu} = 1$ and respectively $\frac{\rho}{\theta} = 1$ ,$\frac{\rho}{\theta} = 0.5$,$\frac{\rho}{\theta} = 0.2$ and $\frac{\rho}{\theta} = 0.1$. Estimated demographic history are represented for all tested germination rate, $\beta = 1$ (red), 0.5 (blue), 0.2 (green) and 0.1 (purple). The demographic history is estimated using a) eSMC where $\beta*$ equal the estimated germination rate, b) MSMC, c) MSMC2 and d) PSMC'.

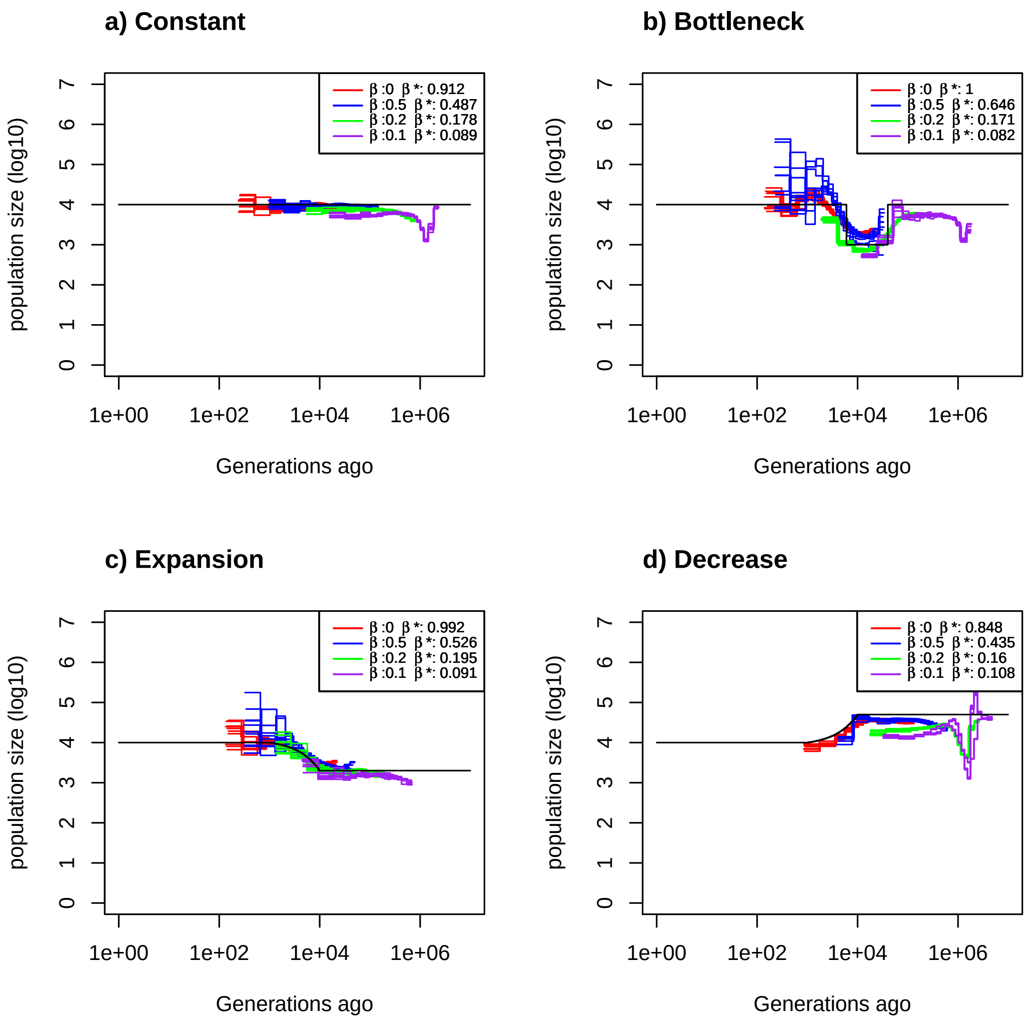


Figure 10: Estimated demographic history using four simulated sequences of 10Mb under four different demographic scenarios with 10 replicates. Mutation and recombination rate are set to $2.5 \times 10^{-8}$ per generation per bp. Simulation were done under four different germination rate $\beta$. We have $\beta = 1$ (red), 0.5 (blue), 0.2 (green) and 0.1 (purple). Therefore $\frac{r}{\mu} = 1$ and we respectively have $\frac{\rho}{\theta} = 1$ ,$\frac{\rho}{\theta} = 0.5$,$\frac{\rho}{\theta} = 0.2$ and $\frac{\rho}{\theta} = 0.1$. The simulated demographic history is represented in black. a) Demographic history simulated under a constant population size. b) Demographic history simulated under a bottleneck. c) Demographic history simulated under an expansion. d) Demographic history simulated under a decrease. In addition we simulated data under four different germination rate $\beta$. $\beta$* equal the estimated germination rate.

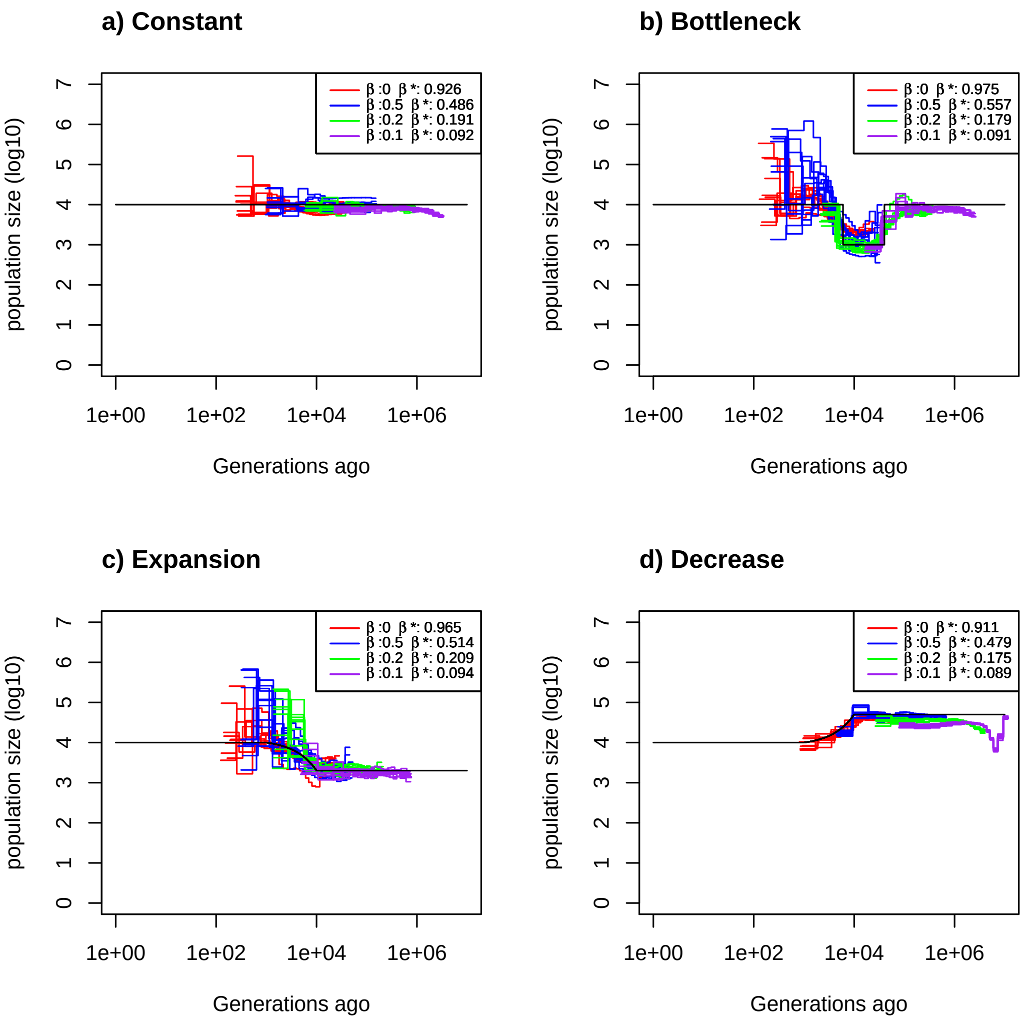


Figure 11: Estimated demographic history using four simulated sequences of 10Mb under four different demographic scenarios with 10 replicates. Mutation and recombination rate are set to $5 \times 10^{-9}$ per generation per bp. Simulation were done under four different germination rate b. We have $\beta = 1$ (red), 0.5 (blue), 0.2 (green) and 0.1 (purple).. Therefore $\frac{r}{\mu} = 1$ and we respectively have $\frac{\rho}{\theta} = 1$ ,$\frac{\rho}{\theta} = 0.5$,$\frac{\rho}{\theta} = 0.2$ and $\frac{\rho}{\theta} = 0.1$. The simulated demographic history is represented in black. a) Demographic history simulated under a constant population size. b) Demographic history simulated under a bottleneck. c) Demographic history simulated under an expansion. d) Demographic history simulated under a decrease. In addition we simulated data under four different germination rate $\beta$. $\beta*$ equal the estimated germination rate.

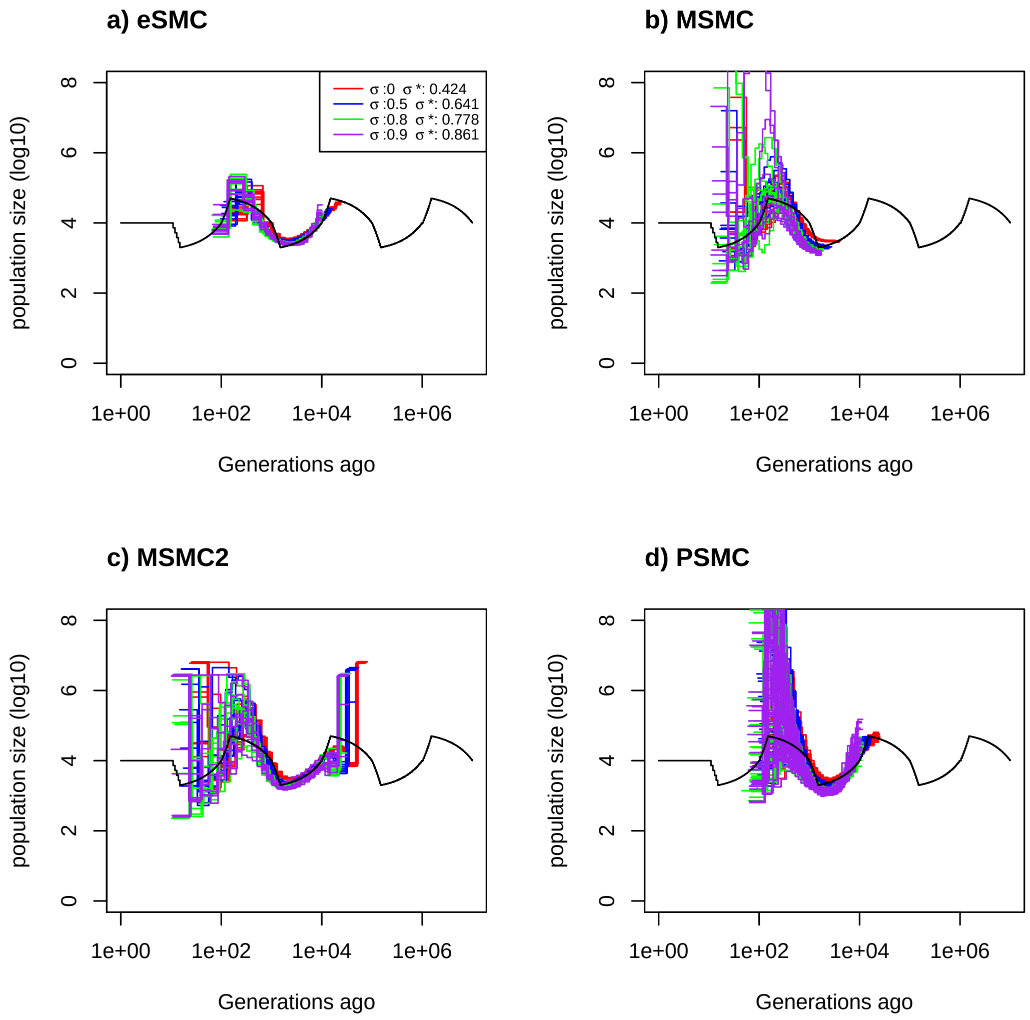


Figure 12: Estimated demographic history using four simulated sequences of 10Mb and ten replicates under a saw-tooth demographic scenario (black). Simulation were done under four different self-fertilization rate $\sigma$ (0,0.5,0.8 and 0.9). The mutation is set to $2.5 \times 10^{-8}$ and the recombination rate to $1.25 \times 10^{-7}$ per generation per bp. Therefore $\frac{r}{\mu} = 5$ and respectively $\frac{\rho}{\theta} = 5$ ,$\frac{\rho}{\theta} = 2.5$,$\frac{\rho}{\theta} = 1$ and $\frac{\rho}{\theta} = 0.5$. Estimated demographic history are represented for all tested self-fertilization, $\sigma = 1$ (red), 0.5 (blue), 0.2 (green) and 0.1 (purple). The demographic history is estimated using a) eSMC where $\sigma*$ equals the estimated self-fertilization rate, b) MSMC, c) MSMC2 and d) PSMC'.

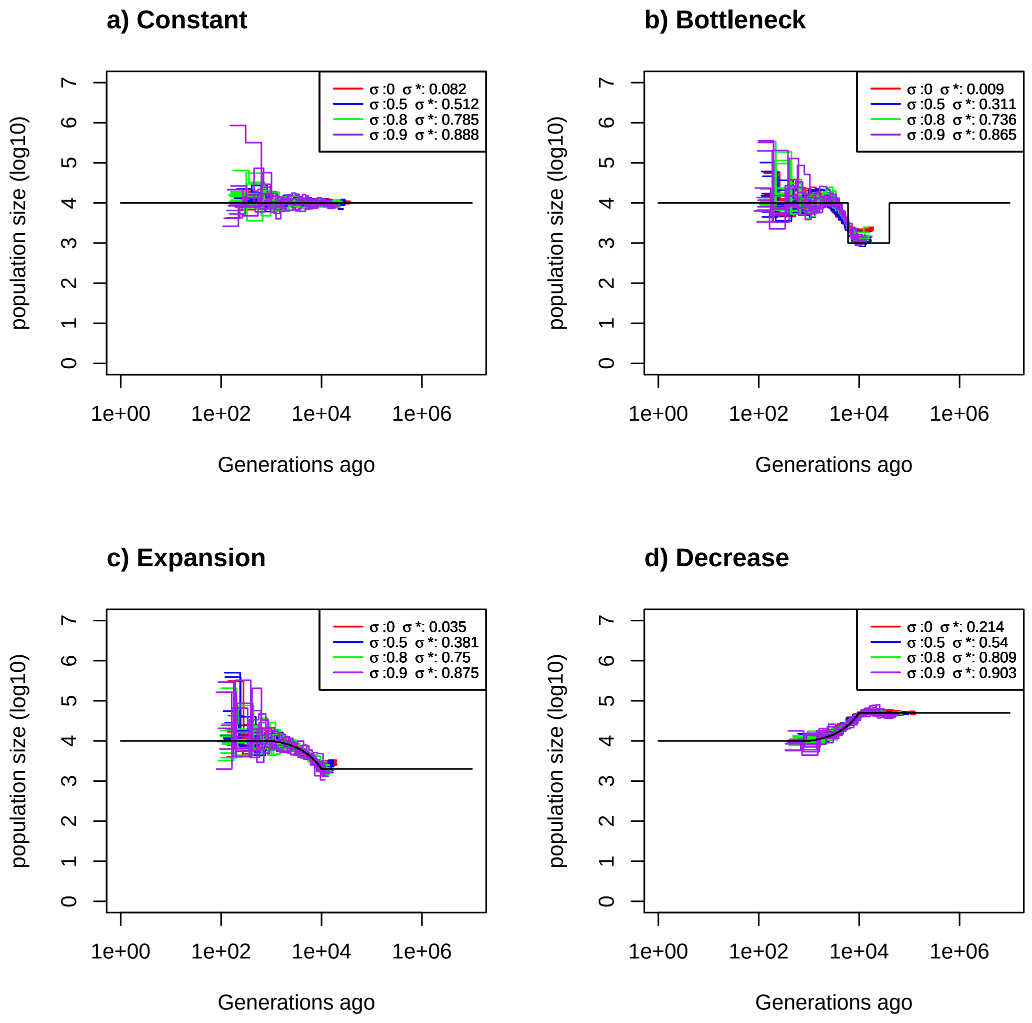


Figure 13: Estimated demographic history using four simulated sequences of 10Mb under four different demographic scenarios with 10 replicates. Mutation and recombination rate are set to $2.5 \times 10^{-8}$ per generation per bp. Simulation were done under four different self-fertilization rate $\sigma$ (0,0.5,0.8 and 0.9). Therefore $\frac{r}{\mu} = 1$ and respectively $\frac{\rho}{\theta} = 1$ ,$\frac{\rho}{\theta} = 0.5$,$\frac{\rho}{\theta} = 0.2$ and $\frac{\rho}{\theta} = 0.1$. The simulated demographic history is represented in black. a) Demographic history simulated under a constant population size. b) Demographic history simulated under a bottleneck. c) Demographic history simulated under an expansion. d) Demographic history simulated under a decrease. In addition we simulated data under four different self-fertilization rate $\sigma$ . We have $\sigma = 0$ (red), 0.5 (blue), 0.8 (green) and 0.9 (purple). $\sigma*$ equal the estimated self-fertilization rate.

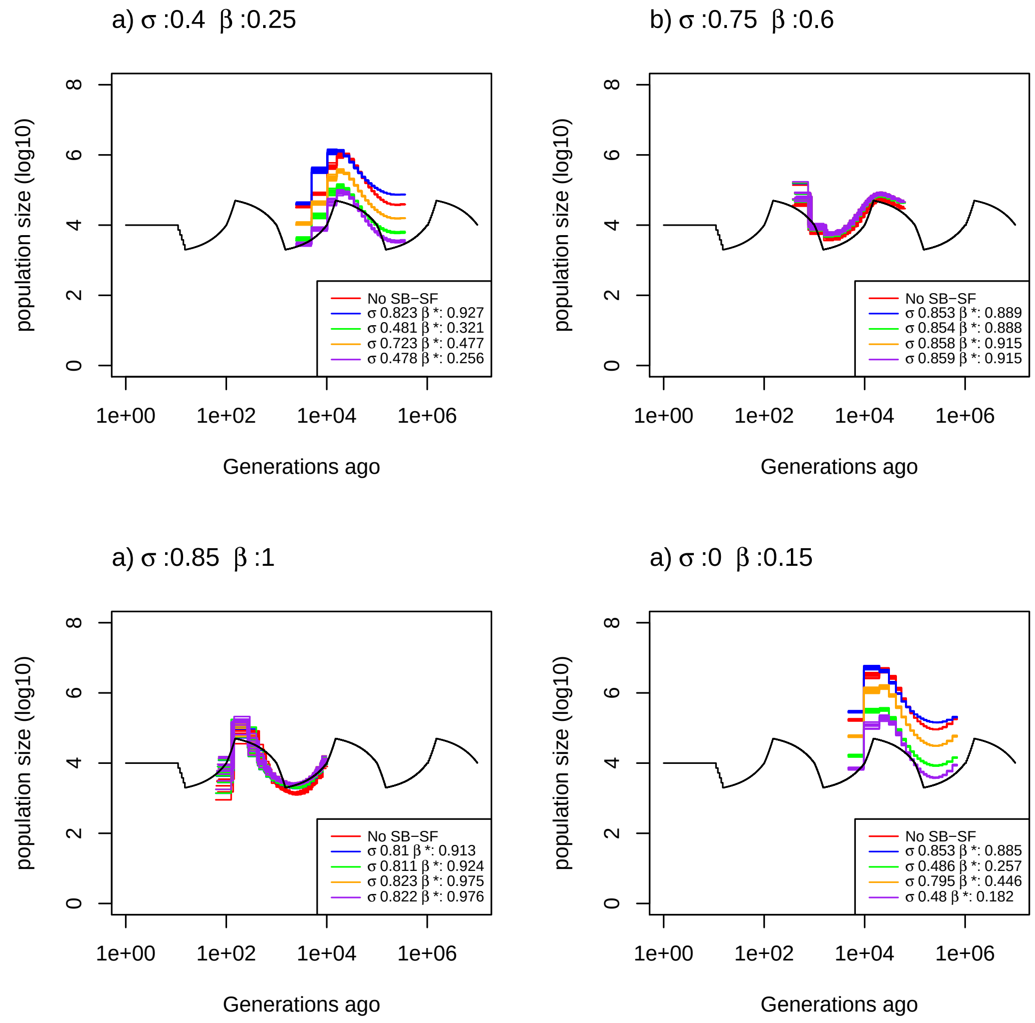


Figure 14: Demographic history estimated by eSMC for ten replicates using four simulated sequences of 10Mb under a saw-tooth demographic scenario and four different combinations of germination (b) and self-fertilization (s) rate but resulting in the same $\frac{\rho}{\theta} = 1$. Mutation rate is set to $2.5 \times 10^{-8}$ and recombination rate to $1.667 \times 10^{-7}$ per generation per bp. Therefore $\frac{r}{\mu} = 6.67$. The four combination are : a) $\sigma = 0.4$ and $\beta = 0.25$, b) $\sigma = 0.75$ and $\beta = 0.6$, c) $\sigma = 0.85$ and $\beta = 1$ and d) $\sigma = 0$ and $\beta = 0.15$. Hence, for each scenario $\frac{\rho}{\theta} = 1$ For each combination of $\beta$ and $\sigma$, eSMC was launched with five different prior settings: ignoring seed banks and self-fertilization (red), accounting for seed banks and self-fertilization but without setting priors (blue), accounting for seed banks and self-fertilization with a prior set only for the self-fertilization rate (green), only for the germination rate (orange) or for both (purple). $\sigma*$ and $\beta*$ respectively represent the estimated self-fertilization and germination rate.

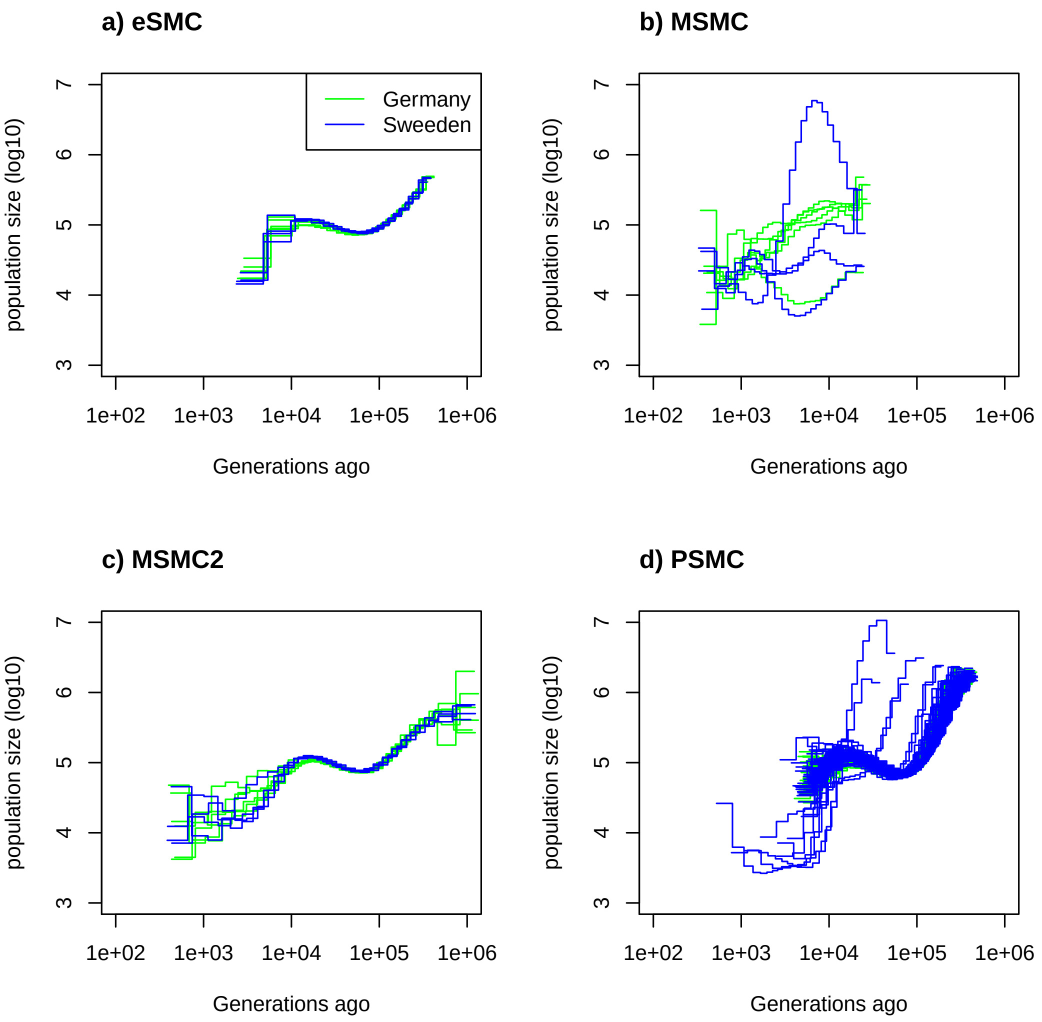


Figure 15: Demographic history of two European (Sweden (blue) and German (green)) populations of *A. thaliana*. Mutation rate is set to $7 \times 10^{-9}$ per generation per bp and was use as prior for recombination rate. a) Demographic history estimated by eSMC without accounting self-fertilzation or dormancy. b) Demographic history estimated by MSMC. c) Demographic history estimated by MSMC2 . d) Demographic history estimated by PSMC'.
