## Appendix Mathematical derivations for "Inference of past demography, dormancy and self-fertilization rates from whole genome sequence data"

### 1 eSMC

To define our Hidden Markov Model (HMM) we need to define :

- Hidden States
- The signal (observed data)
- A Transition matrix (Probability of jumping from one state to another)
- An Emission matrix (Probability of observing the data given the hidden state)
- An Initial probability (Probability of hidden states at the first position of the sequence)

#### 1.1 Notations and Assumptions

We here define the different notations used and their meaning:

- $\beta$ : the germination rate (expected probability to germinate at every generation, between 0 and 1)
- $\sigma$ : self fertilization rate (between 0 and 1)
- $N_0$ : Population at present time
- $r$ : recombination rate per nucleotide per $4N_0$ generations
- $\mu$ : Mutation rate per nucleotide per $4N_0$ generations
- u : recombination time (which follows a continuous uniform distribution on the coalescent tree)
- L: Sequence length in bp
- $\rho = r(L-1)$
- $\theta = \mu L$
- $N_t$ : Population size at time t
- $\chi_t$ : Scaled population size at time t ($N_t = \chi_t N_0$)

The model's assumptions are :

- Piecewise constant population size

- Infinite site model
- Constant mutation, recombination, germination and self-fertilization rate in time
- Constant mutation and recombination rate along the sequence
- Neutrality
- Wright-Fisher model

#### 1.2 Hidden States

We define our hidden states at one position as the coalescent time between the two individual at that position. We note that coalescent time t (t>0). A transition from a coalescent time s to time t ($t \neq s$) at the next can only occur if a recombination happened in between the two positions.

#### 1.3 Observations

Our observation, or the signal, is a sequence of 1 and 0. This sequence is build from phasing the DNA sequences of two individual. When going along the sequence, if both nucleotide are similar, then the signal is 0 (no mutation occurred). If both are different, then a mutation occurred, and the signal is 1 (Figure 1).

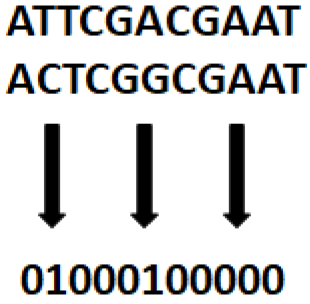


Figure 1: Building the observed data from phased sequences

#### 1.4 Transition Matrix

A transition to state t from state s (t $\neq$ s) can only occur if there is a recombination event. Assuming Recombination event along the sequence as a Poisson process we have the recombination probability of :

$$P(rec|s) = (1 - e^{-\frac{\beta 2(1-\sigma)}{2-\sigma} 2rs}) \tag{1}$$

We now Assume that a recombination event occurred at time u (<s) where u follows a uniform distribution between 0 and s. Then three scenarios are possible. Either the new coalescent time is smaller (t<s),bigger (t>s) or unchanged (t=s). Those scenarios are display on Figure 2 A),C) and B).

### 1.4.1 t<s

The resulting floating branch of the recombination event coalesces at time t < s . This mean it must not coalesce before time t (including itself). In addition we have u<t. The transition probability is therefore :

$$P(t|s,u) = \frac{2\beta^2}{(2-\sigma)\chi_t}(e^{\int_u^t -\frac{4\beta^2}{(2-\sigma)\chi_v} dv}) \tag{2}$$

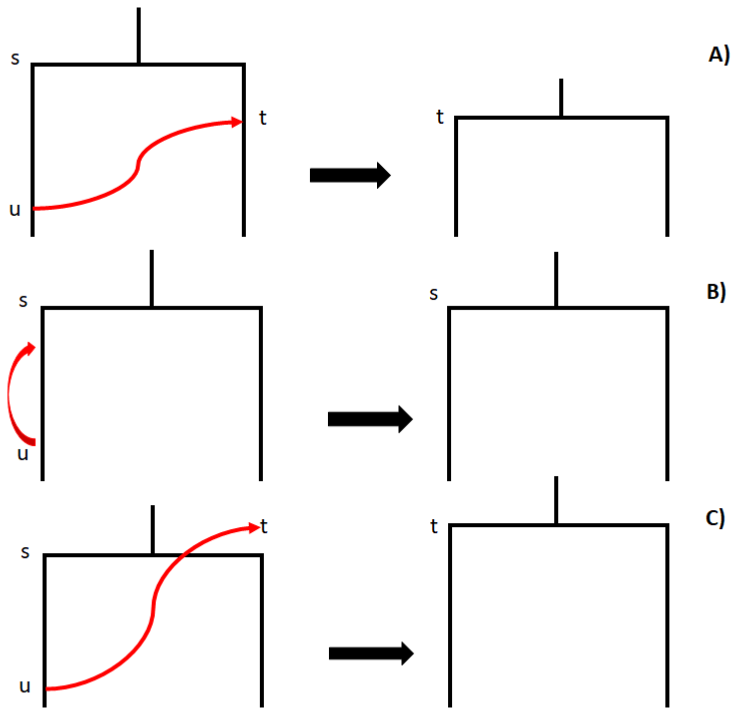


Figure 2: Three possibilities after a recombination event

### 1.4.2 t=s

The resulting floating branch of the recombination event self coalesce before time t. We therefore have the transition probability :

$$P(s|s,u) = \int_u^s \frac{2\beta^2}{(2-\sigma)\chi_k} e^{\int_u^k -\frac{4\beta^2}{(2-\sigma)\chi_v}dv} dk \tag{3}$$

### 1.4.3 t>s

The resulting floating branch of the recombination event must not coalesce (including itself) before time s. Then no coalescent event must happen before time t. We therefore have the transition probability :

$$P(s|s,u) = \frac{2\beta^2}{(2-\sigma)\chi_t} e^{\int_u^s -\frac{4\beta^2}{(2-\sigma)\chi_v}dv} e^{\int_s^t -\frac{2\beta^2}{(2-\sigma)\chi_v}dv} \tag{4}$$

##### 1.4.4 Transition probability in continuous time

In the end we have :

$$p(t|s,u) = \begin{cases} (1-e^{-\frac{\beta 2(1-\sigma)}{2-\sigma}2rs})\frac{2\beta^2}{(2-\sigma)\chi_t}(e^{\int_u^t -\frac{4\beta^2}{(2-\sigma)\chi_v}dv}) & if \quad u<t<s \\ e^{-\frac{\beta 2(1-\sigma)}{2-\sigma}2rs} + (1-e^{-\frac{\beta 2(1-\sigma)}{2-\sigma}2rs})\int_u^s \frac{2\beta^2}{(2-\sigma)\chi_k} e^{\int_u^k -\frac{4\beta^2}{(2-\sigma)\chi_v}dv} dk & if \quad t=s \\ (1-e^{-\frac{\beta 2(1-\sigma)}{2-\sigma}2rs})\frac{2\beta^2}{(2-\sigma)\chi_t} e^{\int_u^s -\frac{4\beta^2}{(2-\sigma)\chi_v}dv} e^{\int_s^t -\frac{2\beta^2}{(2-\sigma)\chi_v}dv} & if \quad t>s \\ 0 & if \quad otherwise \end{cases} \tag{5}$$

Once again,if $\beta = 1$ and $\sigma = 0$ we fall back on the probability from PSMC'.
One can find $p(t|s)$ using the total probability formula which is:

$$p(t|s) = \int_0^s \frac{1}{s} p(t|s,u)du \tag{6}$$

As explained before, the state space must be finite. We therefore discretized time in n intervals. At one point the hidden state is $\alpha$ if $t \in [T_\alpha, T_{\alpha+1}]$, where $\alpha \in [0, (n-1)]$. We define $T_\alpha$ :

$$T_\alpha = -\frac{(2-\sigma)}{2\beta^2}\ln(1-\frac{\alpha}{n}) \tag{7}$$

We therefore have:

$$p(\alpha|s) = \int_{T_\alpha}^{T_{\alpha+1}} p(t|s)dt \tag{8}$$

The transition matrix need to be the probability from one state to another. Therefore we need the probability when the coalescent time at the previous position (which is here s) belongs to the state $\gamma$. To do this we simply replace s by the average coalescent time $t_\gamma$.

##### 1.4.5 Initial Probability

We use the equilibrium probability as initial probability. The equilibrium probability is the probability that the first coalescent happens in each time interval and is thus given by :

$$\begin{aligned}
q_o(\alpha) &= \int_{T_\alpha}^{T_{\alpha+1}} \frac{2\beta^2}{(2-\sigma)\chi_\alpha} e^{\int_0^t \frac{-2\beta^2}{(2-\sigma)\chi_v}dv} dt \\
q_o(\alpha) &= \int_{T_\alpha}^{T_{\alpha+1}} \frac{2\beta^2}{(2-\sigma)\chi_\alpha} e^{\int_0^{T_\alpha} \frac{-2\beta^2}{(2-\sigma)\chi_v}dv} e^{\int_{T_\alpha}^t \frac{-2\beta^2}{(2-\sigma)\chi_v}dv} dt \\
q_o(\alpha) &= e^{\int_0^{T_\alpha} \frac{-2\beta^2}{(2-\sigma)\chi_v}dv} \int_{T_\alpha}^{T_{\alpha+1}} \frac{2\beta^2}{(2-\sigma)\chi_\alpha} e^{\frac{-2\beta^2(t-T_\alpha)}{(2-\sigma)\chi_\alpha}} dt \\
q_o(\alpha) &= e^{\sum_{\eta=0}^{\alpha-1} \frac{-2\beta^2}{(2-\sigma)\chi_\eta}\Delta_\eta}(1-e^{\frac{-2\beta^2\Delta_\alpha}{(2-\sigma)\chi_\alpha}})
\end{aligned} \tag{9}$$

##### 1.4.6 Calculation of $t_\gamma$

$$\begin{aligned}
t_\gamma = E[\text{Coalescent time}|\gamma] &= \frac{E[\text{Coalescent time} \cap \gamma]}{P(\gamma)} = \frac{\int_{T_\gamma}^{T_{\gamma+1}} t\Lambda_\gamma e^{-\int_0^t \Lambda_v dv} dt}{q_0(\gamma)} \\
&= \frac{\Lambda_\gamma \int_{T_\gamma}^{T_{\gamma+1}} t e^{-\int_0^{T_\gamma} \Lambda_v dv} e^{-\int_{T_\gamma}^t \Lambda_v dv} dt}{q_0(\gamma)} = \frac{\Lambda_\gamma e^{-\int_0^{T_\gamma} \Lambda_v dv} \int_{T_\gamma}^{T_{\gamma+1}} t e^{-\int_{T_\gamma}^t \Lambda_v dv} dt}{q_0(\gamma)} \\
&= \frac{\Lambda_\gamma \int_{T_\gamma}^{T_{\gamma+1}} t e^{(T_\gamma - t)\Lambda_\gamma} dt}{(1-e^{-\Delta_\gamma\Lambda_\gamma})} = \frac{T_\gamma - T_{\gamma+1}e^{-\Delta_\gamma\Lambda_\gamma}}{(1-e^{-\Delta_\gamma\Lambda_\gamma})} + \frac{\int_{T_\gamma}^{T_{\gamma+1}} e^{(T_\gamma - t)\Lambda_\gamma} dt}{(1-e^{-\Delta_\gamma\Lambda_\gamma})} \\
&= \frac{T_\gamma - T_{\gamma+1}e^{-\Delta_\gamma\Lambda_\gamma}}{(1-e^{-\Delta_\gamma\Lambda_\gamma})} + \frac{(1-e^{-\Delta_\gamma\Lambda_\gamma})}{\Lambda_\gamma(1-e^{-\Delta_\gamma\Lambda_\gamma})} = \frac{T_\gamma - T_{\gamma+1}e^{-\Delta_\gamma\Lambda_\gamma}}{(1-e^{-\Delta_\gamma\Lambda_\gamma})} + \frac{1}{\Lambda_\gamma}
\end{aligned} \tag{10}$$

Where :

$$\begin{aligned}
\Delta_\gamma &= T_{\gamma+1} - T_\gamma \\
\Lambda_\gamma &= \frac{2\beta^2}{(2-\sigma)\chi_\gamma}
\end{aligned} \tag{11}$$

##### 1.4.7 Calculation of $p(\alpha|\gamma)$

$\alpha < \gamma$ We first need $p(t|t_\gamma)$ when $\alpha < \gamma$, which is obtained as described below :

$$
\begin{aligned}
p(t|t_\gamma) &= \int_0^t \frac{P_\gamma}{t_\gamma} \frac{2\beta^2}{(2-\sigma)\chi_t} (e^{\int_u^t -\frac{4\beta^2}{(2-\sigma)\chi_v} dv}) du \\
&= \frac{P_\gamma}{t_\gamma} \int_0^t \frac{2\beta^2}{(2-\sigma)\chi_t} (e^{\int_u^t -\frac{4\beta^2}{(2-\sigma)\chi_v} dv}) du \\
&= \frac{P_\gamma}{t_\gamma} (\sum_{\eta=0}^{\alpha-1} \int_{T_\eta}^{T_{\eta+1}} \frac{2\beta^2}{(2-\sigma)\chi_t} (e^{\int_u^t -\frac{4\beta^2}{(2-\sigma)\chi_v} dv}) du + \int_{T_\alpha}^t \frac{2\beta^2}{(2-\sigma)\chi_t} (e^{\int_u^t -\frac{4\beta^2}{(2-\sigma)\chi_v} dv}) du) \\
&= \frac{P_\gamma}{t_\gamma} (\sum_{\eta=0}^{\alpha-1} \int_{T_\eta}^{T_{\eta+1}} \frac{2\beta^2}{(2-\sigma)\chi_t} (e^{\int_u^{T_{\eta+1}} -\frac{4\beta^2}{(2-\sigma)\chi_v} dv}) (e^{\int_{T_{\eta+1}}^t -\frac{4\beta^2}{(2-\sigma)\chi_v} dv}) du + \int_{T_\alpha}^t \frac{2\beta^2}{(2-\sigma)\chi_t} (e^{-\frac{(t-u)4\beta^2}{(2-\sigma)\chi_\alpha}}) du) \\
&= \frac{P_\gamma}{t_\gamma} \frac{2\beta^2}{(2-\sigma)\chi_\alpha} (\sum_{\eta=0}^{\alpha-1} e^{\int_{T_{\eta+1}}^t -\frac{4\beta^2}{(2-\sigma)\chi_v} dv} \int_{T_\eta}^{T_{\eta+1}} (e^{-(T_{\eta+1}-u)\frac{4\beta^2}{(2-\sigma)\chi_\eta}}) du + \int_{T_\alpha}^t e^{-\frac{(t-u)4\beta^2}{(2-\sigma)\chi_\alpha}} du) \\
&= \frac{P_\gamma 2\beta^2}{t_\gamma (2-\sigma)\chi_\alpha} (\sum_{\eta=1}^{\alpha-1} \frac{e^{-\int_{T_{\eta+1}}^t \frac{4\beta^2}{(2-\sigma)\chi_v} dv} (1 - e^{-\Delta_\eta \frac{4\beta^2}{(2-\sigma)\chi_\eta}})}{\frac{4\beta^2}{(2-\sigma)\chi_\eta}} + \frac{(1 - e^{(T_\alpha - t)\frac{4\beta^2}{(2-\sigma)\chi_\alpha}})}{\frac{4\beta^2}{(2-\sigma)\chi_\alpha}})
\end{aligned}
\tag{12}
$$

Where:

$$
P_\gamma = (1 - e^{-2rt_\gamma \frac{\beta 2(1-\sigma)}{(2-\sigma)}})
\tag{13}
$$

We can now calculate $p(\alpha|\gamma)$.

$$
\begin{aligned}
p(\alpha|\gamma) &= P_\gamma \int_{T_\alpha}^{T_{\alpha+1}} p(t|t_\gamma) dt \\
&= P_\gamma \int_{T_\alpha}^{T_{\alpha+1}} \frac{2\beta^2}{t_\gamma(2-\sigma)\chi_\alpha} (\sum_{\eta=1}^{\alpha-1} \frac{e^{-\int_{T_{\eta+1}}^t \frac{4\beta^2}{(2-\sigma)\chi_v} dv} (1 - e^{-\Delta_\eta \frac{4\beta^2}{(2-\sigma)\chi_\eta}})}{\frac{4\beta^2}{(2-\sigma)\chi_\eta}} + \frac{(1 - e^{(T_\alpha - t)\frac{4\beta^2}{(2-\sigma)\chi_\alpha}})}{\frac{4\beta^2}{(2-\sigma)\chi_\alpha}}) dt \\
&= P_\gamma \int_{T_\alpha}^{T_{\alpha+1}} \frac{2\beta^2}{t_\gamma(2-\sigma)\chi_\alpha} (\sum_{\eta=1}^{\alpha-1} \frac{e^{-\int_{T_\alpha}^t \frac{4\beta^2}{(2-\sigma)\chi_v} dv} e^{-\int_{T_{\eta+1}}^{T_\alpha} \frac{4\beta^2}{(2-\sigma)\chi_v} dv} (1 - e^{-\Delta_\eta \frac{4\beta^2}{(2-\sigma)\chi_\eta}})}{\frac{4\beta^2}{(2-\sigma)\chi_\eta}} + \frac{(1 - e^{(T_\alpha - t)\frac{4\beta^2}{(2-\sigma)\chi_\alpha}})}{\frac{4\beta^2}{(2-\sigma)\chi_\alpha}}) dt \\
&= \frac{P_\gamma 2\beta^2}{t_\gamma(2-\sigma)\chi_\alpha} (\int_{T_\alpha}^{T_{\alpha+1}} \sum_{\eta=1}^{\alpha-1} \frac{e^{(T_\alpha - t)\frac{4\beta^2}{(2-\sigma)\chi_\alpha} dv} e^{-\int_{T_{\eta+1}}^{T_\alpha} \frac{4\beta^2}{(2-\sigma)\chi_v} dv} (1 - e^{-\Delta_\eta \frac{4\beta^2}{(2-\sigma)\chi_\eta}})}{\frac{4\beta^2}{(2-\sigma)\chi_\eta}} dt + \frac{\Delta_\alpha - \frac{(1-e^{-\Delta_\alpha \frac{4\beta^2}{(2-\sigma)\chi_\alpha}})}{\frac{4\beta^2}{(2-\sigma)\chi_\alpha}}}{\frac{4\beta^2}{(2-\sigma)\chi_\alpha}}) \\
&= \frac{P_\gamma 2\beta^2}{t_\gamma(2-\sigma)\chi_\alpha} (\sum_{\eta=1}^{\alpha-1} \frac{(1 - e^{-\Delta_\alpha \frac{4\beta^2}{(2-\sigma)\chi_\alpha}}) e^{-\int_{T_{\eta+1}}^{T_\alpha} \frac{4\beta^2}{(2-\sigma)\chi_v} dv} (1 - e^{-\Delta_\eta \frac{4\beta^2}{(2-\sigma)\chi_\eta}})}{\frac{4\beta^2}{(2-\sigma)\chi_\alpha} \frac{4\beta^2}{(2-\sigma)\chi_\eta}} + \frac{\Delta_\alpha - \frac{(1-e^{-\Delta_\alpha \frac{4\beta^2}{(2-\sigma)\chi_\alpha}})}{\frac{4\beta^2}{(2-\sigma)\chi_\alpha)}}}{\frac{4\beta^2}{(2-\sigma)\chi_\alpha}}) \\
&= \frac{P_\gamma}{t_\gamma 2} (\sum_{\eta=1}^{\alpha-1} \frac{(1 - e^{-\Delta_\alpha \frac{4\beta^2}{(2-\sigma)\chi_\alpha}}) e^{-\int_{T_{\eta+1}}^{T_\alpha} \frac{4\beta^2}{(2-\sigma)\chi_v} dv} (1 - e^{-\Delta_\eta \frac{4\beta^2}{(2-\sigma)\chi_\eta}})}{\frac{4\beta^2}{(2-\sigma)\chi_\eta}} + \Delta_\alpha - \frac{(1 - e^{-\Delta_\alpha \frac{4\beta^2}{(2-\sigma)\chi_\alpha}})}{\frac{4\beta^2}{(2-\sigma)\chi_\alpha}}) \\
&= \frac{P_\gamma}{t_\gamma 2} (\sum_{\eta=1}^{\alpha-1} \frac{(1 - e^{-\Delta_\alpha \frac{4\beta^2}{(2-\sigma)\chi_\alpha}}) e^{-\sum_{\zeta=\eta+1}^{\alpha} \frac{4\Delta_\zeta \beta^2}{(2-\sigma)\chi_\zeta}} (1 - e^{-\Delta_\eta \frac{4\beta^2}{(2-\sigma)\chi_\eta}})}{\frac{4\beta^2}{(2-\sigma)\chi_\eta}} + \Delta_\alpha - \frac{(1 - e^{-\Delta_\alpha \frac{4\beta^2}{(2-\sigma)\chi_\alpha}})}{\frac{4\beta^2}{(2-\sigma)\chi_\alpha}})
\end{aligned}
\tag{14}
$$

Where:

$$
P_\gamma = (1 - e^{-2rt_\gamma \frac{\beta 2(1-\sigma)}{(2-\sigma)}})
\tag{15}
$$

$\alpha > \gamma$ We first need $p(t|t_\gamma)$ when $\alpha > \gamma$, which is obtained as described below :

$$
\begin{aligned}
p(t|t_\gamma) &= \int_0^{t_\gamma} \frac{P_\gamma 2\beta^2}{t_\gamma(2-\sigma)\chi_t}(e^{\int_u^{t_\gamma} -\frac{4\beta^2}{(2-\sigma)\chi v}dv} e^{\int_{t_\gamma}^{t} -\frac{2\beta^2}{(2-\sigma)\chi v}dv})du \\
&= \frac{P_\gamma 2\beta^2}{t_\gamma(2-\sigma)\chi_\alpha} e^{\int_{t_\gamma}^{t} -\frac{2\beta^2}{(2-\sigma)\chi v}dv} \int_0^{t_\gamma} (e^{\int_u^{t_\gamma} -\frac{4\beta^2}{(2-\sigma)\chi v}dv})du \\
&= \frac{P_\gamma 2\beta^2}{t_\gamma(2-\sigma)\chi_\alpha} e^{\int_{t_\gamma}^{t} -\frac{2\beta^2}{(2-\sigma)\chi v}dv} (\sum_{\eta=0}^{\gamma-1} \int_{T_\eta}^{T_{\eta+1}} (e^{\int_u^{t_\gamma} -\frac{4\beta^2}{(2-\sigma)\chi v}dv})du + \int_{T_\gamma}^{t_\gamma} (e^{\int_u^{t_\gamma} -\frac{4\beta^2}{(2-\sigma)\chi v}dv})du) \\
&= \frac{P_\gamma e^{\int_{t_\gamma}^{t} -\frac{2\beta^2}{(2-\sigma)\chi v}dv} 2\beta^2}{t_\gamma(2-\sigma)\chi_\alpha} (\sum_{\eta=0}^{\gamma-1} \int_{T_\eta}^{T_{\eta+1}} (e^{-(T_{\eta+1}-u)\frac{4\beta^2}{(2-\sigma)\chi v}} e^{\int_{T_{\eta+1}}^{t_\gamma} -\frac{4\beta^2}{(2-\sigma)\chi v}dv})du + \int_{T_\gamma}^{t_\gamma} (e^{-(t_\gamma-u)\frac{4\beta^2}{(2-\sigma)\chi_\gamma}})du) \\
&= \frac{P_\gamma 2\beta^2}{t_\gamma(2-\sigma)\chi_\alpha} e^{\int_{t_\gamma}^{t} -\frac{2\beta^2}{(2-\sigma)\chi v}dv} (\sum_{\eta=1}^{\gamma-1} e^{-\int_{T_{\eta+1}}^{t_\gamma} \frac{4\beta^2}{(2-\sigma)\chi v}dv} \frac{(1-e^{-\Delta_\eta \frac{4\beta^2}{(2-\sigma)\chi_\eta}})}{\frac{4\beta^2}{(2-\sigma)\chi_\eta}} + \frac{(1-e^{(T_\gamma - t_\gamma)\frac{4\beta^2}{(2-\sigma)\chi_\gamma}})}{\frac{4\beta^2}{(2-\sigma)\chi_\gamma}}))
\end{aligned}
\tag{16}
$$

Where:

$$
P_\gamma = (1 - e^{-2rt_\gamma \frac{\beta 2(1-\sigma)}{(2-\sigma)}}) \tag{17}
$$

We can now calculate $p(\alpha|\gamma)$.

$$
\begin{aligned}
q(\alpha|\gamma) &= P_\gamma \int_{T_\alpha}^{T_{\alpha+1}} q(t|t_\gamma)dt \\
&= \int_{T_\alpha}^{T_{\alpha+1}} \frac{P_\gamma 2\beta^2}{t_\gamma(2-\sigma)\chi_\alpha} e^{-\int_{t_\gamma}^{t} \frac{2\beta^2}{(2-\sigma)\chi v}dv} (\sum_{\eta=1}^{\gamma-1} e^{-\int_{T_{\eta+1}}^{t_\gamma} \frac{4\beta^2}{(2-\sigma)\chi v}dv} \frac{(1-e^{-\Delta_\eta \frac{4\beta^2}{(2-\sigma)\chi_\eta}})}{\frac{4\beta^2}{(2-\sigma)\chi_\eta}} + \frac{(1-e^{\frac{(T_\gamma - t_\gamma)4\beta^2}{(2-\sigma)\chi_\gamma}})}{\frac{4\beta^2}{(2-\sigma)\chi_\gamma}})dt \\
&= \frac{P_\gamma 2\beta^2}{t_\gamma(2-\sigma)\chi_\alpha} (\sum_{\eta=0}^{\gamma-1} e^{-\int_{T_{\eta+1}}^{t_\gamma} \frac{4\beta^2}{(2-\sigma)\chi v}dv} \frac{(1-e^{-\Delta_\eta \frac{4\beta^2}{(2-\sigma)\chi_\eta}})}{\frac{4\beta^2}{(2-\sigma)\chi_\eta}} + \frac{(1-e^{\frac{(T_\gamma - t_\gamma)4\beta^2}{(2-\sigma)\chi_\gamma}})}{\frac{4\beta^2}{(2-\sigma)\chi_\gamma}}) \\
&\qquad \int_{T_\alpha}^{T_{\alpha+1}} e^{-\int_{t_\gamma}^{T_\alpha} \frac{2\beta^2}{(2-\sigma)\chi v}dv} e^{-\int_{T_\alpha}^{t} \frac{2\beta^2}{(2-\sigma)\chi v}dv} dt \\
&= \frac{P_\gamma 2\beta^2}{t_\gamma(2-\sigma)\chi_\alpha} (\sum_{\eta=1}^{\gamma-1} e^{-\int_{T_{\eta+1}}^{t_\gamma} \frac{4\beta^2}{(2-\sigma)\chi v}dv} \frac{(1-e^{-\Delta_\eta \frac{4\beta^2}{(2-\sigma)\chi_\eta}})}{\frac{4\beta^2}{(2-\sigma)\chi_\eta}} + \frac{(1-e^{\frac{(T_\gamma - t_\gamma)4\beta^2}{(2-\sigma)\chi_\gamma}})}{\frac{4\beta^2}{(2-\sigma)\chi_\gamma}}) \\
&\qquad e^{-\int_{t_\gamma}^{T_\alpha} \frac{2\beta^2}{(2-\sigma)\chi v}dv} \int_{T_\alpha}^{T_{\alpha+1}} e^{(T_\alpha - t)\frac{2\beta^2}{(2-\sigma)\chi_\alpha}} dt \\
&= \frac{P_\gamma 2\beta^2}{t_\gamma(2-\sigma)\chi_\alpha} (\sum_{\eta=1}^{\gamma-1} e^{-\int_{T_{\eta+1}}^{t_\gamma} \frac{4\beta^2}{(2-\sigma)\chi v}dv} \frac{(1-e^{-\Delta_\eta \frac{4\beta^2}{(2-\sigma)\chi_\eta}})}{\frac{4\beta^2}{(2-\sigma)\chi_\eta}} + \frac{(1-e^{\frac{(T_\gamma - t_\gamma)4\beta^2}{(2-\sigma)\chi_\gamma}})}{\frac{4\beta^2}{(2-\sigma)\chi_\gamma}}) \\
&\qquad e^{-\int_{t_\gamma}^{T_\alpha} \frac{2\beta^2}{(2-\sigma)\chi v}dv} \frac{(1-e^{-\Delta_\alpha \frac{2\beta^2}{(2-\sigma)\chi_\alpha}})}{\frac{2\beta^2}{(2-\sigma)\chi_\alpha}} \\
&= \frac{P_\gamma}{t_\gamma} (\sum_{\eta=1}^{\gamma-1} e^{-\int_{T_{\eta+1}}^{t_\gamma} \frac{4\beta^2}{(2-\sigma)\chi v}dv} \frac{(1-e^{-\Delta_\eta \frac{4\beta^2}{(2-\sigma)\chi_\eta}})}{\frac{4\beta^2}{(2-\sigma)\chi_\eta}} + \frac{(1-e^{\frac{(T_\gamma - t_\gamma)4\beta^2}{(2-\sigma)\chi_\gamma}})}{\frac{4\beta^2}{(2-\sigma)\chi_\gamma}}) e^{-\int_{t_\gamma}^{T_\alpha} \frac{2\beta^2}{(2-\sigma)\chi v}dv} (1-e^{-\Delta_\alpha \frac{2\beta^2}{(2-\sigma)\chi_\alpha}})
\end{aligned}
\tag{18}
$$

Where:

$$
P_\gamma = (1 - e^{-2rt_\gamma \frac{\beta 2(1-\sigma)}{(2-\sigma)}}) \tag{19}
$$

$\alpha = \gamma$ Because probabilities sum up to one. We have the following formula:

$$p(\gamma|\gamma) = 1 - (\sum_{\alpha=0}^{\gamma-1} p(\alpha|\gamma) + \sum_{\alpha=\gamma+1}^{n} p(\alpha|\gamma)) \tag{20}$$

#### 1.5 Emission Matrix

Because of seed banking,the coalescent tree can be very big. In this case the infinite site model hypothesis might be violated,therefore we have the following formula:

$$\begin{aligned} P(0|\gamma) &= e^{-2\mu t_\gamma} \\ P(1|\gamma) &= 1 - e^{-2\mu t\gamma} \end{aligned} \tag{21}$$

Where $\mu$ is the mutation rate per nucleotide per N generation and $t\gamma$ the average coalescent time in state $\gamma$.

#### 1.6 Calculating the objective function of the Baum-Welch Algorithm

To calculate the objective function (CL) we first need to define it. We define it as :

$$CL = P(Y, X|\beta, \chi, \rho) \tag{22}$$

Which is the probability of the signal (Y) and the sequence of Hidden states (X) given the germination rate ($\beta$),self-fertilization rate ($\sigma$),recombination rate ($\rho$) and population size per interval ($\chi$). To calculate this probability we use a forward-backward algorithm.

##### 1.6.1 Forward Algorithm

The Forward algorithm is an iterative algorithm that calculate at step t the probability :

$$fo_t(i) = P(Y_{1,\ldots,t}, X_t = i) \tag{23}$$

To calculate this probability we define:

- T : Transition matrix ($T_{i,j} = P(X(t) = j|X(t-1) = i)$)
- O : observation matrix ($O_{i,i} = P(Y(t) = e(t)|X(t) = i)$) where e(t) is the observed data at position t ( which can be 0 or 1)

**Initialization** $fo_1 = q_0 O_1$

Where $q_0$ is the vector of initial probabilities.

**Recursive formula** $fo_t = O_t T^T fo_{t-1}$

In case of recurrent pattern in the sequence, a technique has been developed to accelerate the algorithm[1]. Example, if there are many repetition of the same observation (repetition of length l), we then have :

$$fo_t = O_t T^T fo_{t-1} = O_t T^T O_{t-1} T^T fo_{t-2} = (O_t T^T)^l fo_{t-l} \tag{24}$$

To compute $P(O_{1,\ldots,L})$ which we call the likelihood (LH), We simply notice that :

$$\sum_i fo_t(i) = \sum_i P(Y_{1,\ldots,t}, X_t = i) = P(Y_{1,\ldots,t}) \tag{25}$$

Which lead to :

$$\begin{aligned} c_t &= \sum_i fo_t(i) \\ fo_t^* &= \frac{fo_t}{c_t} \\ fo_t &= O_t T^T fo_{t-1}^* \\ LH &= \prod_{t=1}^{L} c_t \end{aligned} \tag{26}$$

##### 1.6.2 Backward Algorithm

The backward algorithm is an iterative algorithm that calculates : $ba_t(i) = P(Y_{t+1,\dots,L}|X_t = i)$
The notations are the same as before. The algorithm is defined as :

**Initialization** $ba_L = I$

**Recursive formula** $ba_{t-1} = TO_t ba_t$

In a similar way, if there are repeated observations in the sequence we have:

$$ba_t = TO_{t+1}ba_{t+1} = TO_{t+1}TO_{t+2}ba_{t+2} = (TO_t)^l ba_{t+l} \tag{27}$$

##### 1.6.3 Baum-Welch Algorithm

**The classic algorithm** The Baum-Welch Algorithm is a particular case of the generalized Expectation-Maximization algorithm (Biology Sequence Analysis R. Durbin, Inference in Hidden Markov Models O. Cappé ).At every step the algorithm update the parameters that maximize the function Q. Where Q is defined at step t as :

$$Q(\theta|\theta^t) = \sum_X P(X|Y, \theta^t) log(P(X, Y|\theta)) \tag{28}$$

And so :

$$\theta^{t+1} = argmax_\theta Q(\theta|\theta^t) \tag{29}$$

We have :

$$Q(\theta|\theta^t) = \sum_X P(X|Y, \theta^t) log(\prod_{X_1} P(X_1|\theta)^{N(X_1)} \prod_{X,Z} P(X|Z, \theta)^{N(X,Z)} \prod_{X,Y} P(Y|X, \theta^t)^{N(Y,X)}) \tag{30}$$

Where :

- $N(X_1)$ : number of first position where the hidden state is $X_1$
- $N(X, Z)$ : number of transition from state Z to X
- $N(Y, X)$ : number of position with observation Y happening with hidden state X

Which gives us :

$$Q(\theta|\theta^t) = \nu_{\theta^t} log(P(X_1|\theta)) + \sum_{X,Y} E(X, Z|\theta^t) log(P(X|Z, \theta)) + \sum_{X,Y} E(Y, X|\theta^t) log(P(Y|X, \theta)) \tag{31}$$

Where:

- $\nu_\theta$ : The equilibrium probability conditional to the set of parameters $\theta$

- $P(X_1|\theta)$ : Probability of the first hidden state conditional to the set of parameters $\theta$
- $E(X,Z|\theta^t)$ : Expected number of transition of X from Z conditional to the observation and set of parameters $\theta^t$
- $P(X|Z,\theta)$ : Transition Probability from state Z to state X conditional to the set of parameters $\theta$
- $E(Y,X|\theta^t)$ Expected number of observation of type Y that happened during state X conditional to the observation and set of parameters $\theta^t$
- $P(Y|X,\theta)$ : Emission probability conditional to the set of parameters $\theta$

However, the objective function used in [2] is

$$Q^*(\theta|\theta^t) = \sum_{X,Z} E(X,Z|\theta^t)log(P(X|Z,\theta)) \tag{32}$$

**Calculating** $E(X,Z|\theta^t)$

$$E(X,Z|\theta^t) = \frac{\sum_{l=1}^{L-1} fo_t(Z)ba_{t+1}(X)P(Y_{t+1}|X)P(X|Z)}{P(Y_{1,\dots,L}|\theta)} \tag{33}$$

**Calculating** $E(Y,X|\theta^t)$

$$E(Y,X|\theta^t) = \frac{\sum_{t=1}^{L} fo_t(X)ba_t(X)1_{Y_t}}{P(Y_{1,\dots,L}|\theta)} \tag{34}$$

##### 1.6.4 Speeding the algorithm

A trick to speed the algorithm can be find in the supplementary material of [3]. The trick is to skip position with no segregating sites. We thus have (Cf forward and backward algorithm)

$$\begin{aligned} fo_l &= (W^T)^{l-k} fo_k \\ ba_k &= W^{l-k} ba_l \\ W &= TO = PDP^{-1} \end{aligned} \tag{35}$$

**Calculating** $E(Y,X|\theta^t)$ We want $fo_l ba_l$. We thus have:

$$\begin{aligned} E(Y,X|\theta^t) = \sum_{i=k}^{l-1} fo_i ba_i &= \sum_{i=0}^{l-k-1} diag((W^T)^i fo_k (W^{l-k-i} ba_l)^T) \\ \sum_{i=k}^{l-1} fo_i ba_i &= \sum_{i=0}^{l-k-1} diag((W^T)^i fo_k ba_l^t (W^{l-k-i})^T) \\ \sum_{i=k}^{l-1} fo_i ba_i &= \sum_{i=0}^{l-k-1} diag(((P^{-1})D^i P^T fo_k ba_l^t (P^{-1})^T D^{l-k-i} P^T) \\ \sum_{i=k}^{l-1} fo_i ba_i &= diag((P^{-1})^T A P^T) \\ A &= \sum_{i=0}^{l-k-1} D^i P^T fo_k ba_l^t (P^{-1})^T D^{l-k-i} \\ U &= P^T fo_k ba_l^t (P^{-1})^T \\ A &= \sum_{i=0}^{l-k-1} D^i U D^{l-k-i} \end{aligned} \tag{36}$$

We have :

$$\sum_{i=0}^{l-k-1} D^i U D^{l-k-i} = \sum_{i=0}^{m} D^i U D^{m-i} D$$
$$(\sum_{i=0}^{l-k-1} D^i U D^{l-k-i})_{ab} = \sum_{i=0}^{m} D^i_{aa} U_{ab} D^{m+1-i}_{bb} = U_{ab} \sum_{i=0}^{m} D^i_{aa} D^{m+1-i}_{bb} \tag{37}$$

We therefore define Q:

$$Q_{ab} = \sum_{i=0}^{m} D^i_{aa} D^{m-i}_{bb} \tag{38}$$

In the end:

$$A = (U * Q)D \tag{39}$$

Where $*$ stands for the Hadamard product.

**Calculating** $E(X, Z|\theta^t)$ In a similar way.

$$E(X, Z|\theta^t) = \sum_{i=k}^{l-1} \xi_i = (fo_i(ba_{i+1} Y_{i+1})) * T$$
$$\xi_{i,ab} = P(X_i = b, X_{i+1} = a|Y, \theta^t) \tag{40}$$

Which gives us:

$$\sum_{i=k}^{l-1} \xi_i = \sum_{i=k}^{l-1} (fo_i ba^T_{i+1} O) * T$$
$$\xi_{i,ab} = P(X_i = b, X_{i+1} = a|Y_{1,\ldots,L}, \theta^t) \tag{41}$$

With repeated observed data we have:

$$\sum_{i=k}^{l-1} \xi_i = \sum_{i=0}^{l-k-1} ((W^T)^i fo_k ba^T_l (W^{k-l-1-i})^T O) * T$$
$$\sum_{i=k}^{l-1} \xi_i = \sum_{i=0}^{l-k-1} ((P^{-1})^T D^i P^T fo_k ba^T_l (P^{-1})^T D^{l-k-1-i} P^T) O) * T \tag{42}$$
$$\sum_{i=k}^{l-1} \xi_i = (((P^{-1})^T (U * Q) P^T) O) * T$$

#### 1.7 Maximizing the objective function

To maximize the Complete Likelihood, as shown before we need to maximize the following value:

$$Q^*(\theta|\theta^t) = \sum_{X,Z} E(X, Z|\theta^t) log(P(X|Z, \theta)) \tag{43}$$

To maximize the objective function we use a Barzilai-Borwein spectral method.

### References


[1] Andreas Sand, Martin Kristiansen, Christian N. S. Pedersen, and Thomas Mailund. zipHMMlib: a highly optimised HMM library exploiting repetitions in the input to speed up the forward algorithm. *BMC BIOINFORMATICS*, 14, NOV 22 2013.

[2] Stephan Schiffels and Richard Durbin. Inferring human population size and separation history from multiple genome sequences. *NATURE GENETICS*, 46(8):919–925, AUG 2014.

[3] Jonathan Terhorst, John A. Kamm, and Yun S. Song. Robust and scalable inference of population history froth hundreds of unphased whole genomes. *NATURE GENETICS*, 49(2):303–309, FEB 2017.
